# No room to roam: King Cobras reduce movement in agriculture

**DOI:** 10.1101/2020.03.24.006676

**Authors:** Benjamin Michael Marshall, Matt Crane, Inês Silva, Colin Thomas Strine, Max Dolton Jones, Cameron Wesley Hodges, Pongthep Suwanwaree, Taksin Artchawakom, Surachit Waengsothorn, Matt Goode

**Affiliations:** Suranaree University of Technology, Nakhon Ratchasima, Thailand; King Mongkut’s University of Technology Thonburi, Bangkok, Thailand; Population and Community Development Association, Bangkok, Thailand; Sakaerat Environmental Research Station, Nakhon Ratchasima, Thailand; School of Natural Resources and Environment, University of Arizona, Tucson, AZ, USA

**Keywords:** snake, reptile, *Ophiophagus hannah*, elapid, space-use, step-selection, dBBMM, site fidelity, tropical

## Abstract

Studying animal movement provides insights into how animals react to land-use changes, specifically how animals can change their behaviour in agricultural areas. Recent reviews show a tendency for species to reduce movements in response to increased human landscape modification, but the study of movement has not been extensively explored in reptiles. We examined movements of a large reptilian predator, the King Cobra (*Ophiophagus hannah*), in Northeast Thailand. We used a consistent regime of radio-telemetry tracking to document movements across protected forest and adjacent agricultural areas. We then adapted GPS-targeting analytic methods to examine the movement using metrics of site reuse and dynamic Brownian Bridge Movement Model derived motion variance. Examination of motion variance demonstrated that King Cobra movements increased in forested areas and tended to decrease in agricultural areas. Our Integrated Step-Selection Functions indicated that when moving in agricultural areas King Cobras restricted their movements, thereby remaining within vegetated semi-natural areas, often located along the banks of irrigation canals. Site reuse metrics of residency time and number of revisits remained unaffected by distance to landscape features. Neither motion variance nor reuse metrics were consistently affected by the presence of threatening landscape features (e.g. roads, human settlements); suggesting that King Cobras will remain in close proximity to threats, provided habitat patches are available. Although King Cobras displayed heterogeneity in their response to agricultural landscapes, the overall trend suggested a reduction in movements when faced with fragmented habitat patches embedded in an otherwise inhospitable land-use matrix. Reductions in movements are consistent with findings for mammals and forest specialist species.

## Introduction

Examining animal movement can provide important information on conspecific interactions (Jellen et al. 2007), predator-prey dynamics (Courbin et al. 2016, Vogt et al. 2018), reproductive behaviours (Kamath and Losos 2018), and responses to anthropogenic threats (Valeix et al. 2012, Loveridge et al. 2017). Additionally, and perhaps most important to conservation planning, is the connection between movement and resource acquisition (Prange et al. 2004, Mueller et al. 2011, Doherty et al. 2019). Understanding habitat requirements, via animal movement, can help prioritise areas to protect from land-use conversion, inform management, and build conservation plans (Fraser et al. 2018).

Anthropogenic land-use can alter the ecology of a landscape, changing resources (Arrondo et al. 2018), modifying behaviour (Gaynor et al. 2018), and introducing novel threats (Robertson et al. 2013). Such changes can result in increased mortality of species, or even subtler sub-lethal costs (Cottontail et al. 2009, Clark et al. 2011, Karraker et al. 2018). A global review of non-volant mammals revealed that movements are directly impacted by human landscapes: animals present in human landscapes reduce movement (Tucker et al. 2018).

Despite indications of overall reductions in vagility, the impacts of anthropogenic landscapes on threatened species is likely to vary. Evolutionary history and key traits are likely to modify a species’ movements in relation to human-dominated landscapes (Fahrig 2007). For example, species that evolved in continuous habitat (i.e. forest specialists) historically experienced lower costs to large movements and crossing habitat barriers, potentially resulting in species leaving prime habitat and using riskier anthropogenic landscapes (Fahrig 2007).

Vulnerability in anthropogenic landscapes can be augmented by species traits such as large body size, parental investment in offspring, habitat specialisation, and low population densities, which have been connected to increased extinction risk (Purvis et al. 2000, Cardillo 2005, Böhm et al. 2016, Slavenko et al. 2016, Todd et al. 2017). Species frequently involved in human-wildlife conflict are also more vulnerable to direct mortality in anthropogenic landscapes (Shankar et al. 2013, Marshall et al. 2018).

We aimed to explore the movement patterns of a large-bodied, highly persecuted predator in a mixed-use landscape. Reptiles’ role in ecosystems are underappreciated (Miranda 2017) and, in South East Asian agricultural systems, likely constitute an important aspect of the remaining wildlife. Few reptile species fulfil similar ecological functions comparable to large mammals, but King Cobras (*Ophiophagus hannah* [CANTOR, 1836]) share several traits that could indicate their importance in ecosystem functioning and vulnerability to habitat modification. Using radio-telemetry, we 1) assess non-random habitat selection and 2) identify changes in movement patterns within anthropogenic areas to reveal how King Cobras are potentially adapting to land-use change.

## Methods and Materials

### Field Methods

We studied King Cobras at the Sakaerat Biosphere Reserve located in Nakhon Ratchasima province, Northeast Thailand (14.44–14.55°N, 101.88–101.95°E). The reserve is comprised of three zones varying in levels of human-modification: the core zone, protected and fully forested; the buffer zone, protected and undergoing reforestation; and the transitional zone, an agricultural matrix dominated by rice, corn and sugar. The transitional zone also contains 159 villages and a four-lane highway that connects Nakhon Ratchasima to Bangkok. Further descriptions of the study site can be found in Silva et al. (2018) and Marshall et al. (2018, 2019).

The capture and implantation methods, alongside King Cobra measurements, have been previously described in Marshall et al. (2018, 2019). We tracked individuals four times a day, with approximately four hours between tracks from 2014-03-22 to 2018-07-28 (06:30, 11:00, 16:00, 20:00; the distribution of time lags between tracking is available in Supplementary Figure 1). Full details of the tracking protocols can be found in Silva et al. (2018). We named every individual according to their age class, sex and capture number (e.g. AM006 = an adult male who was the sixth King Cobra captured).

### Environmental data

We obtained daily rainfall and temperatures from five weather stations within the Sakaerat Biosphere Reserve core zone to identify seasons (Sakaerat Environmental Research Station 2018). We averaged daily readings by station, and ran cluster analysis to generate seasons using the *segclust2D* package (v.0.2.0 Patin et al. 2018).

For the entire study site, we obtained high quality land-use shapefiles from a land survey by the Thai Land Development Department (Land Development Department, Thailand 2017). We converted categorical land-use classifications to continuous raster layers, describing Euclidean distances to key landscape features (i.e., forest, roads, semi-natural areas, settlements, and water bodies). We set the cell size of the newly created rasters to approximately 10m, which was sufficiently small to detect fine-scale changes. Semi-natural areas were areas of scrub and vegetation not actively being farmed, often along field margins, irrigation canals and in disused plots.

### Motion variance and area estimation

Traditionally, research on reptile spatial ecology has relied on kernel density and minimum convex polygon approaches to estimate space-use, as a proxy for movements and activity. Kernel density estimators are problematic, because the technique assumes independence between locations, which can never be strictly met in radio tracking datasets (Fieberg 2007). Efforts to combat autocorrelation (Worton 1989), lead to a loss of information decreasing the biological relevance of space-use estimates (De Solla et al. 1999). Dynamic Brownian Bridge Movement Models (dBBMM; Kranstauber et al. 2012) present an alternative that accounts for non-independence of locations and provides a balance between over- and under-estimating space-use (Silva et al. 2018, 2020b).

We used the *move* package (v.3.1.0 Kranstauber et al. 2016) to run dBBMMs estimating motion variance and the area used by King Cobras. We used the *adehabitatHR* package to extract utilisation distributions and contours (v.0.4.16 Calenge 2006), and the *rgeos* (v.0.4.2 Bivand and Rundel 2018) package to estimate the area of contours. We used dBBMMs instead of standard BBMMs, because the former allowed for estimates of changes in motion variance over time (Horne et al. 2007, Kranstaubery et al. 2012). Following Kranstauber et al. (2012), we selected a window and margin size for dBBMMs based on a timeframe that was biologically relevant to suspected changes in behavioural states. Due to our reliance on Very High Frequency (VHF) radio tracking and associated coarse temporal resolution data, we targeted the identification of activity and sheltering. We were able to detect shifts from activity to sheltering with slightly greater than one day of radio tracking effort; therefore, we set margin size at 5 data points. A relevant time for a behavioural state to last was approximately one week (i.e. long-term sheltering); therefore, we set window size to 25 data points. We used GPS error for dBBMM location error on a point-by-point basis, for points that did not have GPS error recorded we used the mean GPS error for that individual.

We explored seasonal changes in motion variance and how it was impacted by an individual’s proximity to landscape features (i.e., forest, roads, semi-natural areas, settlements, and water bodies). Due to serial autocorrelation and over dispersal in motion variance and distance raster values, we used non-metric multidimensional scaling (NMDS) to explore interactions among these variables. Using the *vegan* package (v.2.5.5 Oksanen et al. 2019), we ran NMDS on a distance matrix created from the rasters that described distances from key landscape features (using 2000 iterations to produce two axes). We plotted the resulting two-dimensions and coloured points corresponding to the motion variance values. The resulting visualisation allowed us to identify areas of high or low motion variance and the manner in which they are associated with snakes’ distances to landscape features.

### Site fidelity and reuse

Shelter sites are important for species requiring extended periods of low mobility to digest meals (Siers et al. 2018) or undergo ecdysis (Dodd and Barichivich 2007). Reptile studies often infer important areas using the 50% use (“core area”) contour from a kernel density home range estimation. Use of a more intensive radio tracking regime allowed us to identify individual shelter sites, time spent within shelters, and frequency of reuse.

We identified site reuse with the *recurse* package (v.1.1.0 Bracis et al. 2018). We defined each site as a circular area with a radius equal to the mean GPS error recorded for each individual (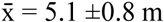, range = 3.5 – 10.0 m). When examining the frequency of revisits, we filtered out sites where the snake was present for less than the mean time between data points (9 hours). We determined whether a site was within the protected core area or the human-modified agricultural areas, then compared how frequency of site reuse and time spent at a shelter differed between these areas. We used Man-Whitney tests (we determined the data was non-normal with qq-plots) to compare differences at a population level.

Time spent at sites (residency time) and the reuse rate (recurse) have direct connections to the extent of animal space use, making them useful metrics to detect restrictions in movement (Van Moorter et al. 2016). To examine these two metrics we ran four Bayesian models in JAGS using the *jagsUI* package (v.1.5.0 Kellner 2018). Two models used a log normal distribution to explore the impacts of proximity to uncorrelated landscape features on log transformed residency time (Bracis et al. 2018). Two models used a Poisson distribution to explore the impacts of proximity to uncorrelated landscape features on revisit counts (Bracis et al. 2018). We determined spatial correlation in the landscape rasters and created two groups of uncorrelated variables (r<0.6) to use as predicators: 1) roads, forest, and settlements; 2) roads, forest, and semi-natural areas.

Because of the variation between individuals, we used the individual ID as a random effect impacting both models’ intercepts and gradients. We excluded AM007 from models because he remained in the forest; therefore, had little opportunity to display preference beyond forests or interact with landscape features.

We used Cauchy and half Cauchy distributions (Lemoine 2019) as hyperparameters for the centre and precision of normal distributions priors for individual random effects on distance to forest, semi-natural areas, roads, settlements and water. We selected weakly informative priors based on the assumption that King Cobras would follow similar movement patterns as those described in Tucker et al. (2018): reduced movement associated with anthropogenic features. We made the prior for the effect of distance to forest negative, reflecting the likely opposite effect from anthropogenic features proximity. We ran all models using three chains over 20,000 interactions, with the first 5,000 discarded as burn-in and a thinned factor of 50. Full JAGS models specifications can be found in at DOI: 10.5281/zenodo.3666029.

We identified convergence via 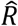 values and traceplots. We evaluated model performance using DIC, Bayes P-values, followed by visual inspection of posterior predictive check plots.

The *recurse* package also allowed us to quantify time spent in the protected core zone of the reserve. Comparing movements to a shapefile of the reserve’s protected zone allowed us to create a summary of all boundary crossings (entrance and exit times). From the revisit data, we calculated overall time spent in the core zone and plotted the use of the zone over time.

### Integrated Step-selection Function

We explored King Cobra movements through the landscape with Integrated Step-Selection Functions (ISSF) from the *amt* package (v.0.0.6 Signer et al. 2018), using the same distance from landscape features rasters used in the above analyses. For ISSF, we inverted raster layers to guard against zero-inflation in distance values and make interpretation of resulting effects more intuitive. We used the landscape values at the endpoints in ISSF, because our sampling regime was temporally insufficient to assume straight-line movements between locations. We produced 200 random locations per step, with no resampling of data, because temporal resolution of our radio-tracking data was coarser than GPS data allowing high numbers of random steps without requiring prohibitively intense computation. Producing 200 random locations reduced the chance of missing rare landscape types, and made the best use of high-resolution raster data (Thurfjell et al. 2014).

All nine models included step length and angle (Forester et al. 2009), with random step lengths and angles drawn from gamma and von Mises distributions, respectively. One model only included step length and angle as predictors, five models included step length, angle and distance from a landscape feature, and three models included step length, angle and a combination of distances from multiple uncorrelated landscape features. We selected models per individual using Aikike’s Information Criterion (AIC), discarding those with a Δ AIC > 2 (Burnham and Anderson 2010). We did not model average to produce a population level model, because we observed high individual heterogeneity. We excluded AM007 from the ISSF analysis, because he never left forested areas.

### Software and data

We completed all analysis in R (v.3.5.3 R Core Team 2019) and R Studio (v.1.2.1335 R Studio Team 2019). The full dataset, with code scripts, can be found at DOI: 10.5281/zenodo.3666029. Movement data is also available on MoveBank (Movebank ID: 1093796277).

For data manipulation, we used R packages *broom* (v.0.5.2 Robinson and Hayes 2019), *data*.*table* (v.1.12.2 Dowle and Srinivasan 2019), *dplyr* (v.0.8.3 Wickham et al. 2019), *forcats* (v.0.4.0 Wickham 2019a), *lubridate* (v.1.7.4 Grolemund and Wickham 2011), *openxlsx* (v.4.1.0 Walker 2018), *readr* (v.1.3.1 Wickham et al. 2018), *reshape2* (v.1.4.3 Wickham 2007), and *stringr* (v.1.4.0 Wickham 2019b). We handled rasters and shapefiles with R packages *raster* (v.2.8.19 Hijmans 2019), *rgdal* (v.1.4.3 Bivand et al. 2019) and *sp* (v.1.3.1 Pebesma and Bivand 2005, Bivand et al. 2013). For visualisations we used R packages *cowplot* (v.0.9.4 Wilke 2019), *ggplot2* (v.3.2.1 Wickham 2009), *ggpubr* (v.0.2 Kassambara 2018), *ggspatial* (v.1.0.3 Dunnington 2018), *scales* (v.1.1.0 Wickham and Seidel 2019) and *scico* (v.1.1.0 Pedersen and Crameri 2018). To determine model convergence and evaluate model performance, we used the R packages *ggmcmc* (v.1.2 Fernández-i-Marín 2016), *ggridges* (v.0.5.1 Wilke 2018), and *tidybayes* (v.1.0.4 Kay 2019).

## Results

We tracked seven King Cobras for an average of 649.7 ±112.3 days (Table 1; all ± indicated the standard error [SE] associated with the 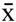, calculated using the *pracma* package (v.2.2.5 Borchers 2019)) enabling us to use dynamic Brownian Bridge Movement Model analysis and explore impacts of land-use on movement. We tracked and located each King Cobra an average of 1834 ±297.1 times, with an average of 8.5 ±0.1 hours between fixes (range = 0.1 – 793.9 hours; Supplementary Figure 1). King Cobras occupied an average of 524 ±104.5 unique locations, covering large areas in protected and unprotected areas (Table 1; Figure 1), with adult males tending to move more. The two juvenile males differed greatly from each other, likely the result of JM013’s northward travel. We only radio tracked a single adult female, which used the smallest area of any snake.

**Figure 1.**
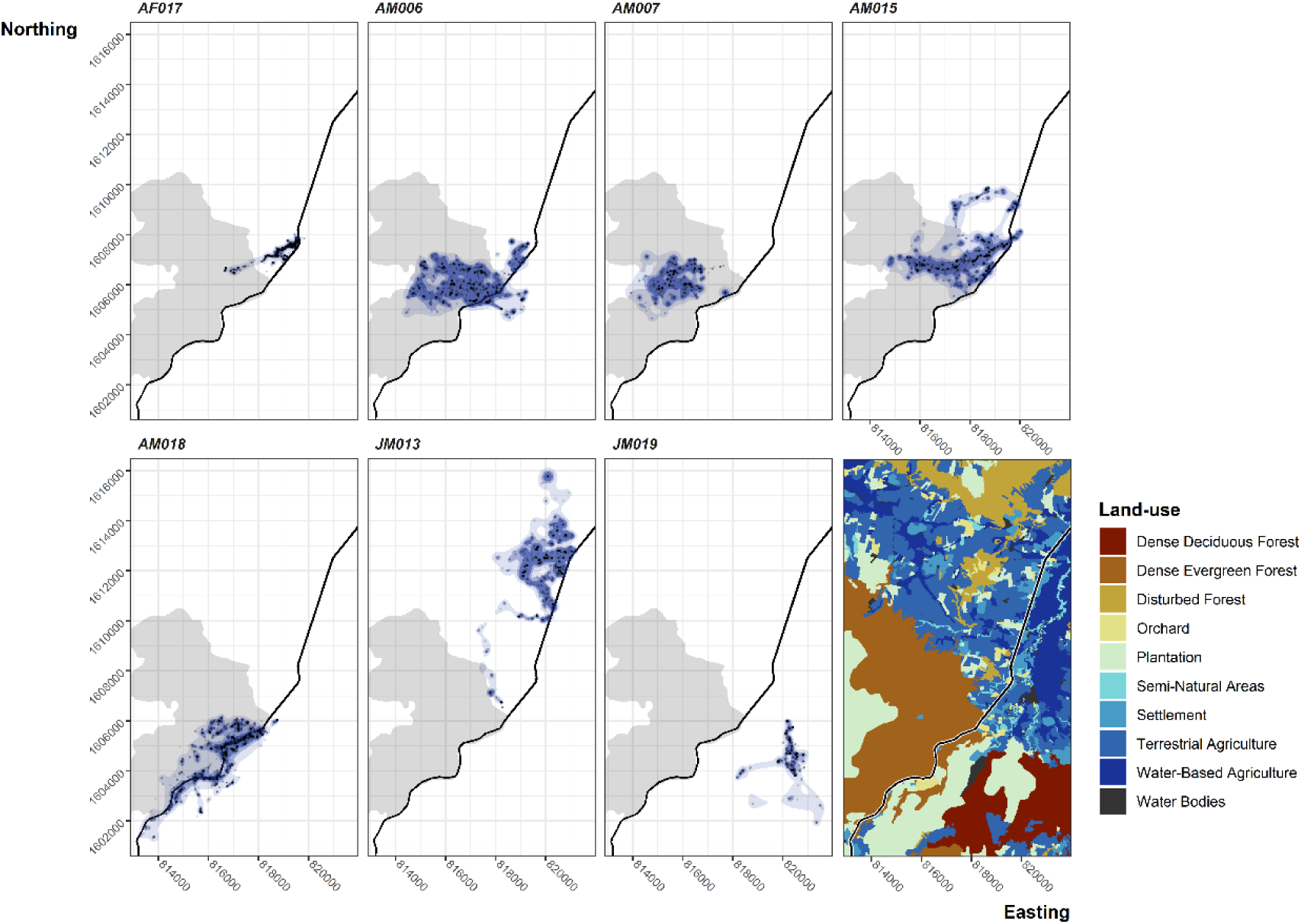
Dynamic Brownian Bridge Movement Model estimates of utilisation distribution contours. Areas displayed with increasing levels of opacity are the 99%, 95% and 90% utilisation contours. Black dots show locations. The shaded background area shows the protected core area. Dark central line is the four-lane 304 highway. Bottom right map shows the land-use types in the area.

**Table 1.** Summary of tracking and movements. Datapoints = number of datapoints collected on an individual irrespective of move or not. Relocations = number of unique locations visited by an individual. Days = number of days tracked. Revisits frequency = the number of days between revisits to a previously used location (days tracked / count of reused locations). Time stationary = mean sheltering time ±SE in days. dBBMM Range = range areas estimated using dBBMM 99%, 95%, and 90% contours. σ^2^m = mean motion variance ±SE. % Outside of PA = Percentage of total time tracked an individual was outside of the protected area.

| ID | Datapoints | Days | Relocations | Revisit frequency | Time stationary | dBBMM Range (ha) | | | $\sigma^2m$ | % outside PA |
| --- | --- | --- | --- | --- | --- | --- | --- | --- | --- | --- |
|  |  |  |  |  |  | 90 | 95 | 99 |  |  |
| AF017 | 2245 | 774.97 | 728 | 3.19 | $1.84 \pm 0.13$ | 41.69 | 68.15 | 149.28 | $7.53 \pm 0.33$ | 91.53 |
| AM006 | 2173 | 723.05 | 542 | 19.03 | $2.58 \pm 0.27$ | 519.60 | 701.44 | 1063.42 | $42.61 \pm 1.74$ | 15.49 |
| AM007 | 969 | 320.66 | 220 | 12.83 | $2.61 \pm 0.53$ | 232.70 | 345.62 | 616.90 | $51.90 \pm 3.81$ | 0.67 |
| AM015 | 1944 | 680.13 | 587 | 13.60 | $2.24 \pm 0.22$ | 379.80 | 603.32 | 1081.54 | $27.3 \pm 1.22$ | 67.57 |
| AM018 | 3122 | 1176.10 | 985 | 7.79 | $2.16 \pm 0.14$ | 255.09 | 492.54 | 977.84 | $33.56 \pm 1.41$ | 42.39 |
| JM013 | 1497 | 561.19 | 381 | 21.58 | $3.09 \pm 0.39$ | 354.33 | 533.26 | 972.74 | $22.35 \pm 1.11$ | 99.95 |
| JM019 | 890 | 311.79 | 228 | 11.99 | $2.56 \pm 0.37$ | 61.01 | 119.04 | 390.39 | $7.90 \pm 0.63$ | – |

Our examination of seasonality using *segclust2D* suggested that five clusters and 23 segments was the best way of dividing the 2012-2018 period into seasons. However, it resulted in seasons unique to single years. Therefore, we manually simplified the seasons into three groups that appear in nearly all years: hot (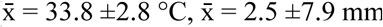 rainfall), wet ((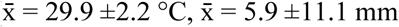 rainfall) and dry (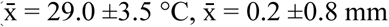 rainfall).

Time spent in human disturbed areas varied dramatically between individuals and showed modest seasonal patterns (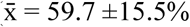, range = 0.7 – 100%; Table 1, Figure 2). During the start of the hot season (Figure 2 red highlight, February-April), adult males ventured out of protected forested areas, a pattern particularly clear in AM006’s movements. During other times of the year, snakes exhibited more consistent use of the protected area, which coincided with more frequent long-term use of shelter sites (Figure 2). The female, AM017, showed a consistent yearly pattern of entering the protected area via a semi-natural area corridor that connected to a streambed.

**Figure 2.**
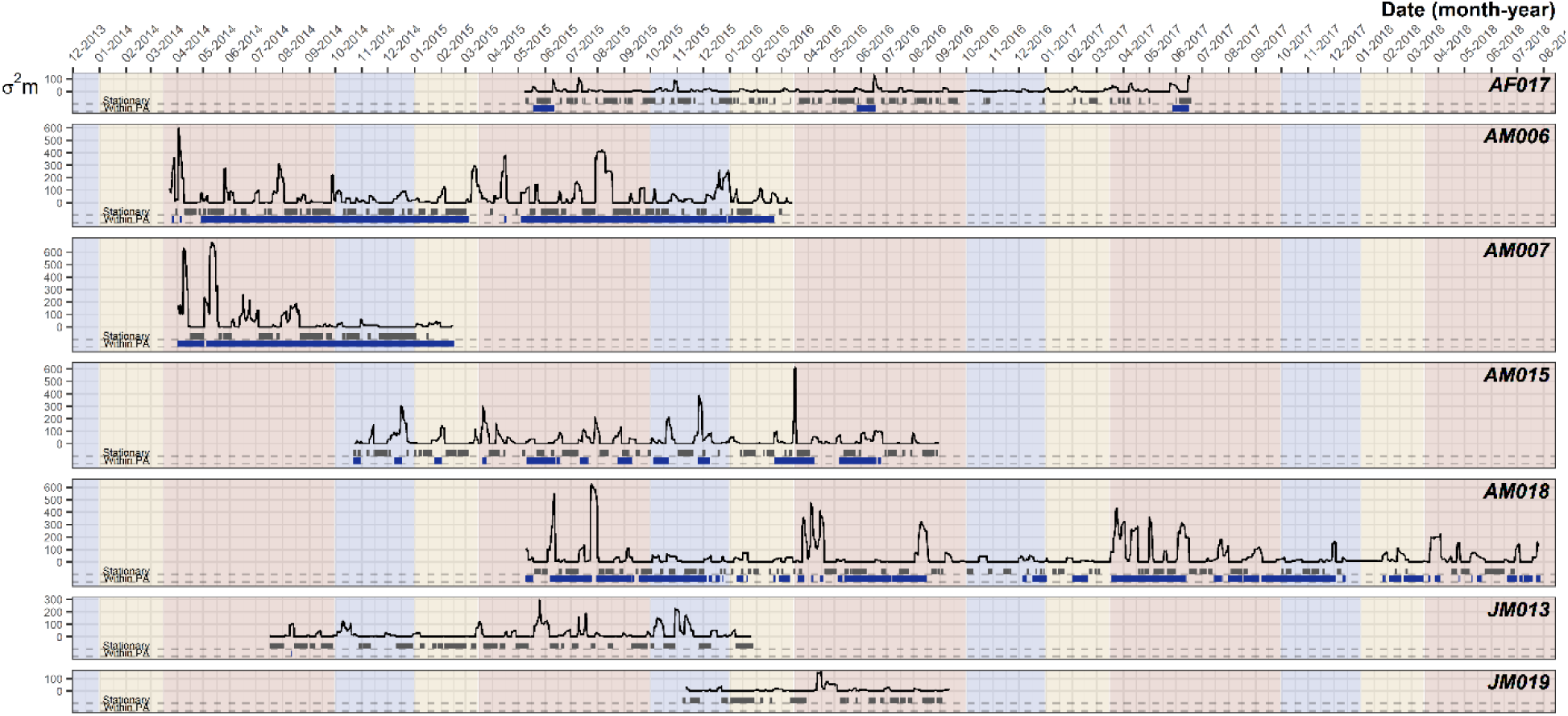
Motion variance of each individual over their tracking period. Black lines show the motion variance values over time. Grey bars indicate long-term sheltering behaviour (i.e., when the time sheltering was greater than the individual’s mean sheltering time). Blue bars indicate times when the individual was within the protected forested area. Shading shows the three seasons: red = Hot, blue = Wet, yellow = Dry.

### Motion variance

Mean motion variance differed among individuals (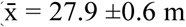, range = 5.6e^-05^ – 675.8 m, Table 1). The largest motion variances belonged to adult males, characterized by larger movements concentrated at the beginning of the hot season (Figure 2, red highlight). Juvenile males did not move as far as adult males at any time of the year, but they did appear to be more active than the female, AF017. Motion variance of AF017 peaked during the hot season, when she entered the protected area in mid-to late-April and left in mid-May. All individuals displayed seasonal differences in motion variance, with the lowest values during the dry season 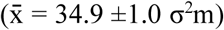 compared to hot 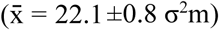 and wet seasons (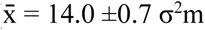; Figure 2).

Motion variance was highest in evergreen and disturbed forests (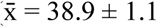, max = 665 m; 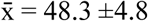, max = 598 m), and lowest in orchards (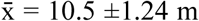, max = 449), semi-natural areas (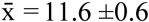, max = 347 m), and water bodies (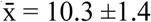, max = 119 m).

Using NMDS, we successfully reduced dimensionality of chosen landscape features, revealing several patterns. The clearest pattern was the grouping of higher motion variance values, the majority of which arose when snakes were < 100 m from forested areas (Figure 3; see Supplementary Figure 2 for bi-plot). In contrast to movement variance values near or within forests, NMDS revealed consistently lower values within 100 m of semi-natural areas. All other covariates were more weakly associated with particular motion variance values. Roads contained a wide array of values, which overlapped with forest and semi-natural areas, suggesting a weaker impact on motion variance. Settlements and water bodies revealed similarly weak associations to motion variance, but there was a tendency for motion variance near or within settlements to be lower than those near or within forests.

**Figure 3.**
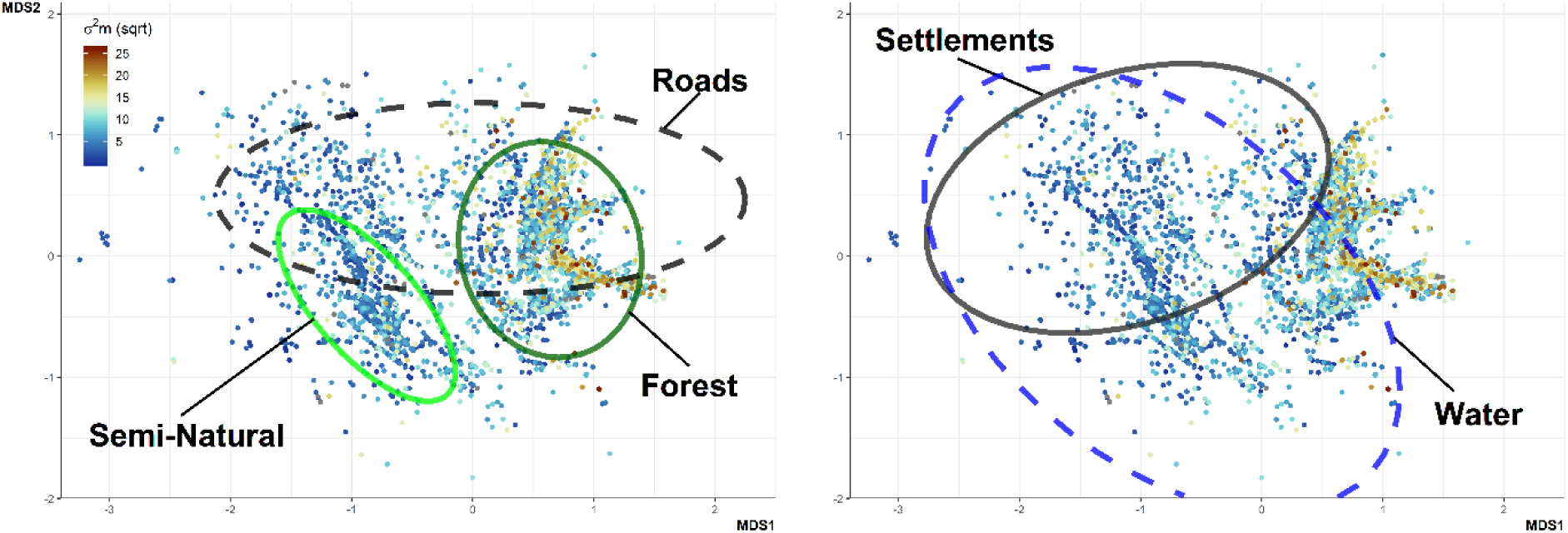
Non-metric multidimensional scaling plot. Motion variance values are reflected by the colour of the points, we have rooted these values so value differences are easier to distinguish. Ellipses indicate 95% of points within 100 m of a given landscape feature. A) Ellipses highlight points existing within 100 m of forest, semi-natural areas, and roads. B) Ellipses highlights points existing within 100 m of water, and settlements.

### Site fidelity and reuse

The *recurse* analysis revealed that mean time spent at a shelter site (stationary for more than 9 hours) was 2.3 ±0.1 days (range = 0.4 – 43.5 days; Supplementary Figure 3), and all snakes demonstrated site fidelity, revisiting a previously used site on average every 13.0 ±2.4 days (range = 3.2 – 21.6 days; Table 1).

Mann-Whitney U tests at a population level failed to detect significant differences in frequency of site reuse between sites in core forest (median = 190.4) and anthropogenic areas (median = 99.1; W = 13027, n_1_ = 99, n_2_ = 250, p-value = 0.4432 two-sided; Supplementary Figure 4). Similarly, time spent within sites was not significantly different between core forest (median = 24.7) and anthropogenic areas (median = 24.8; W = 210410, n_1_ = 398, n_2_ = 1095, p-value = 0.3087 two-sided; Supplementary Figure 5).

All models we ran to predict residency time and revisits converged and produced Bayes p-values close to 0.5. All the models revealed that the distance to landscape features has a negligible effect on residency time or revisit frequency, illustrated by all ß coefficient 95% credible intervals values overlapping with zero. Results of all Bayesian models can be found in Supplementary Table 1.

### Integrated Step-selection Function

Individual movements of the King Cobras were best described by three models (Figure 4; Table 2; full ISSF results can be found in Supplementary Table 2). Model 7 performed best for four individuals, and included proximity to forest, roads, and semi-natural areas. Universally, the locations of AF017, AM015, AM018 and JM019 were positively associated with forests. However, the association between movements, roads, and semi-natural areas varied; AF017, AM015 and JM019 prefered semi-natural areas, but were inconsistently associated with roads. The movements of AM006, AM015, and AF017 while in agricultural land exemplifies King Cobras’ reliance on semi-natural areas (Figure 5). By contrast, AM018’s locations were associated with roads, while weakly avoiding semi-natural areas. But for AM018 model 8 was within 2 Δ AIC. Model 8 replaced semi-natural areas with settlements as a predictor, indicating positive association (ß = 2.504, 95% CI −0.244 – 5.253). Models targeting JM013’s movements were similarly inconclusive, with four models achieving Δ AIC < 2 (including the null model), indicating distance to landscape feature was a poor predictor of movement. Finally, AM006’s movements were best described by model 6, indicating a weak association with water bodies.

**Figure 4.**
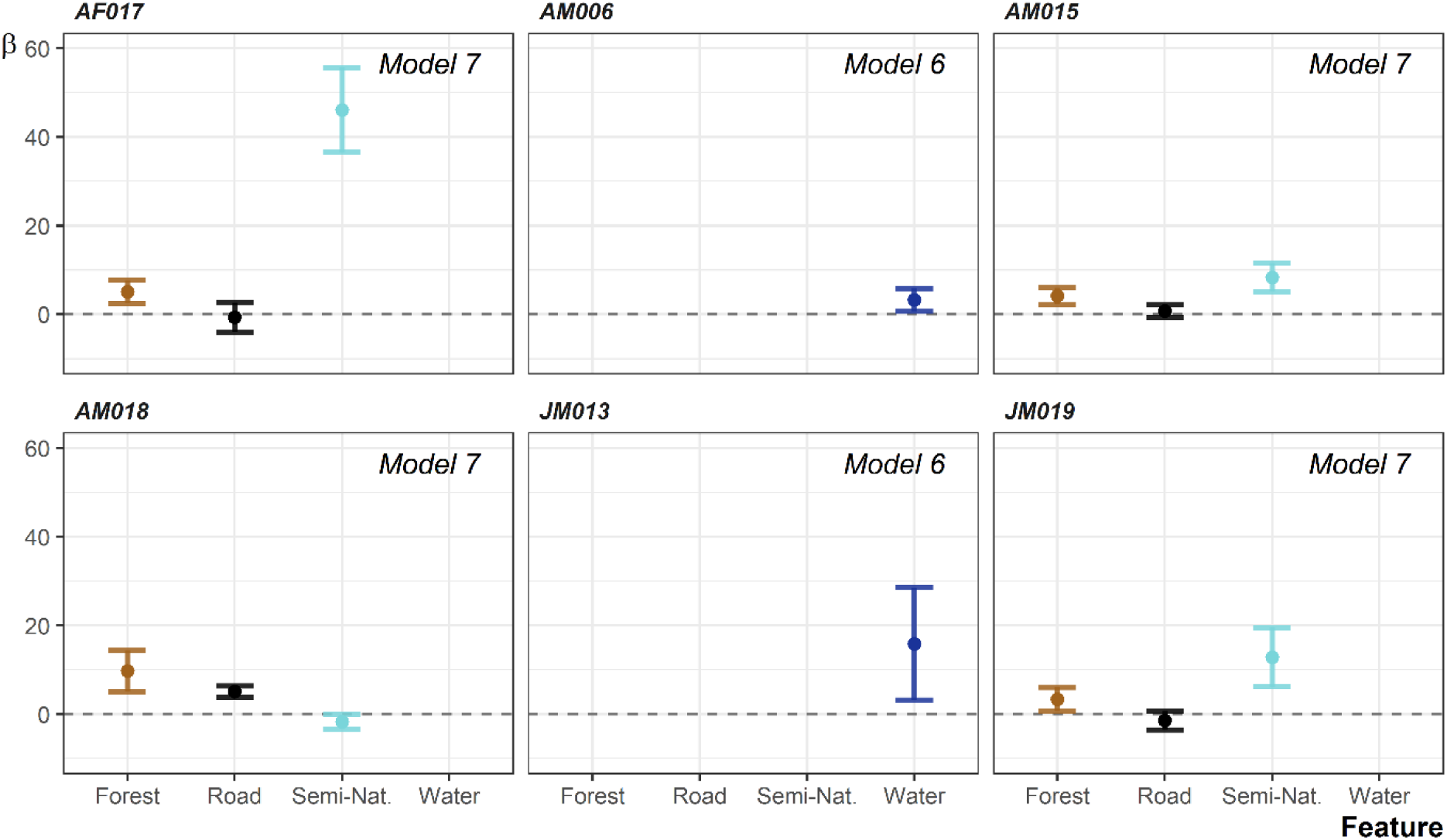
The coefficients from the best performing integrated step-selection functions per individual. Error bars show the 95% confidence interval. JM013 and AM018’s had other models within 2 Δ AIC.

**Figure 5.**
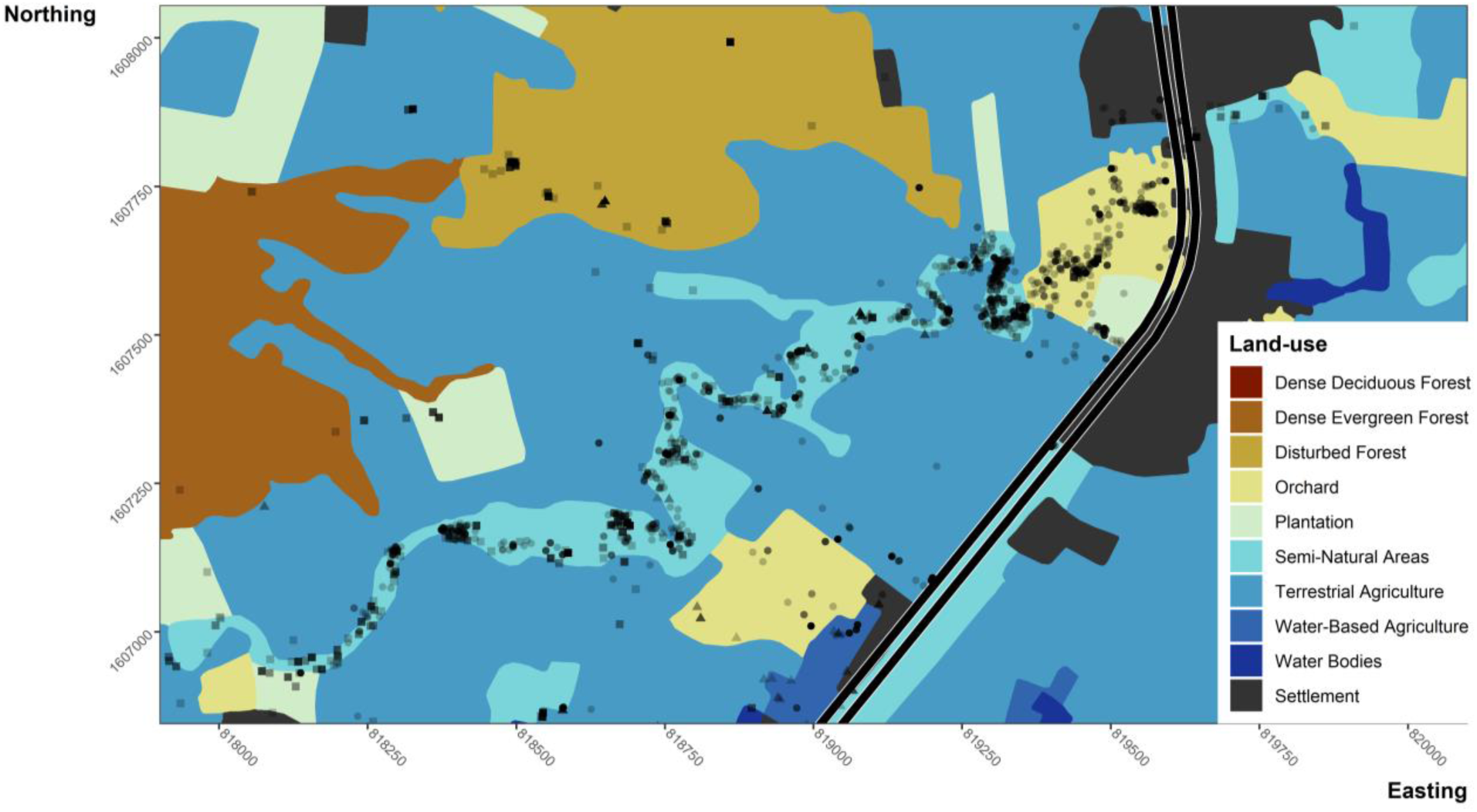
A map of land-use illustrating how King Cobra movements are largely occurring within semi-natural areas. Displayed using semi-transparent points, are the locations of King Cobras across the entire study period. Circles = AF017, triangles = AM006, squares = AM015.

**Table 2.** ISSF model formulation and AIC results. sl = step length; ta = turning angle; dist_* = distance from forest, settlement, semi-natural area, road, and water. * and bold indicate the models < 2 Δ AIC from the top-performing model.

| Model | Model formula, all begin with Model1 formulation | AF017 | AM006 | AM015 | AM018 | JM013 | JM019 |
| --- | --- | --- | --- | --- | --- | --- | --- |
| Model1 | log_sl*cos_ta+strata(step_id_) | 7573.47 | 5794.50 | 6205.87 | 10567.77 | **4014.21\*** | 2402.41 |
| Model2 | Model1+dist_forest+dist_forest:log_sl+dist_forest:cos_ta | 7555.47 | 5794.62 | 6187.03 | 10530.19 | **4013.58\*** | 2399.25 |
| Model3 | Model1+dist_settle+dist_settle:log_sl+dist_settle:cos_ta | 7542.05 | 5794.64 | 6201.81 | 10552.30 | **4014.60\*** | 2402.45 |
| Model4 | Model1+dist_semiNat+dist_semiNat:log_sl+dist_semiNat:cos_ta | 7492.52 | 5783.74 | 6186.73 | 10563.71 | 4017.76 | 2392.32 |
| Model5 | Model1+dist_road+dist_road:log_sl+dist_road:cos_ta | 7557.50 | 5791.24 | 6201.98 | 10498.78 | 4017.31 | 2406.31 |
| Model6 | Model1+dist_water+dist_water:log_sl+dist_water:cos_ta | 7566.34 | **5779.88\*** | 6199.01 | 10559.55 | **4013.23\*** | 2404.11 |
| Model7 | Model1+dist_road+dist_forest+dist_semiNat | **7471.48\*** | 5789.37 | **6171.16\*** | **10471.26\*** | 4017.55 | **2388.28\*** |
| Model8 | Model1+dist_road+dist_forest+dist_settle | 7539.43 | 5789.9 | 6195.03 | **10472.37\*** | 4017.61 | 2400.04 |
| Model9 | Model1+dist_road+dist_forest+dist_water | 7530.33 | 5789.75 | 6187.41 | 10474.02 | 4015.71 | 2398.12 |

## Discussion

We present some of the first evidence for how a large tropical reptile modifies its movements when in agricultural areas. Motion variance was characterized by seasonal peaks associated with breeding activity, but generally showed decreased activity in agricultural areas. Reduced movement in anthropogenic systems reflects meta-analyses of mammalian movements in anthropogenic systems (Tucker et al. 2018). Research on reptile spatial ecology has documented either reduced space-use (Mitrovich et al. 2009, Hoss et al. 2010, Breininger et al. 2011, Lomas et al. 2019) or reduced movement (Parent and Weatherhead 2000, Corey and Doody 2010, Beale et al. 2016, Doherty et al. 2019) in fragmented agricultural landscapes. But the response to fragmentation is not universal, other research failed to detect significant shifts in movement patterns (Row et al. 2012, Wolf et al. 2013, Anguiano and Diffendorfer 2015), or even revealed increased space-use in fragmented areas (Kapfer et al. 2010, Ettling et al. 2013), potentially due to species-specific evolutionary history traits (Fahrig 2007). King Cobras appeared to be reacting in a way consistent with forest specialists, or taxa that have evolved in continuous habitat (Fahrig 2007) –limited boundary avoidance, large movements, and mortality in human-dominated areas (Marshall et al. 2018). Landscape-specialist species occupying fragmented areas likely face limited resources, resulting in restricted movements to more naturalistic corridors (Dondina et al. 2019).

The clearest pattern we documented was preferential use of semi-natural vegetation patches when King Cobras moved within agricultural areas. These patches primarily consist of dense vegetation arrayed linearly along the banks of irrigation canals, and are used more frequently than the surrounding matrix of agricultural fields, acting as movement corridors through the fragmented landscape. Linear habitats potentially impact movements in other reptile species (Kay et al. 2016, Doherty et al. 2019). Doherty et al. (2019) suggested that reduced movement by Eastern Bearded Dragons (*Pogona barbata*) was partly driven by higher prey availability in linear vegetation patches. Although we lack direct evidence suggesting semi-natural areas within agricultural landscapes host relatively higher prey abundance, it is likely King Cobra prey can be found more frequently where vegetation and water are present (Murphy et al. 1999, Barnes et al. 2017, Strine et al. 2018). However, increased movements in forests, at least for some individuals, may indicate that resource availability alone fails to explain variation in movement patterns. Intact forests are extremely valuable and present a resource-rich environment, theoretically reducing the need for foraging movements (Wasko and Sasa 2012, Doherty and Driscoll 2018).

Ectotherms also have to consider the thermal qualities of habitats, shifting habitat usage to maximise efficiency (Blouin-Demers and Weatherhead 2001, 2002). Compared to temperate regions, evidence from the tropics that behavioural shifts are required to maximise thermoregulation is more ambiguous (Luiselli and Akani 2002), but not unknown in larger species (Shine and Madsen 1996). Open fields and vegetation corridors present two contrasting thermal environments. When temperatures are high tropical snakes may need to seek cooler, covered, and environments potentially richer in shelter sites.

Utilisation of covered areas may also be tied to threat avoidance, as threats are known to influence animal movement (Rio-Maior et al. 2019, Suraci et al. 2019). In our study area, roads pose a major threat to many animals (Silva et al. 2020a). King Cobras also fall victim to both direct (Marshall et al. 2018) and indirect (Strine et al. 2014) human-caused mortality. However, we failed to detect clear avoidance of roads or human settlements; King Cobras made use of suitable habitat types regardless of their proximity to threatening landscape features. Similarly, patterns of site reuse remained consistent in respect to proximity to landscape features. This suggests that the overarching driver of site residency time and revisitation is largely independent of habitat, instead likely connected to cycles of ecdysis and prey capture and digestion (Dodd and Barichivich 2007, Siers et al. 2018).

Building on our results, we suggest that future conservation research focus on landscape connectivity. Irrigation canals and forest fragments may allow King Cobras to persist across areas largely separated from protected forest. Research on landscape connectivity could be especially beneficial if paired with an assessment of how threats can be effectively mitigated. The apparent lack of threat avoidance illustrated by the studied King Cobras demands changes in human behaviour. For example, road crossing structures in combination with fencing would likely help to mitigate the threat posed by roads (Rytwinski et al. 2016). Whereas reducing persecution of King Cobras will require a change in current negative perceptions (Shankar et al. 2013, Marshall et al. 2018) and improvements in humane snake removal services, although the cost-effectiveness of snake removal services needs further quantification (Devan-Song et al. 2016).

## Conclusion

Our results indicate that limited areas in agricultural landscapes are suitable for King Cobras, resulting in reduced movements that largely occur within vegetated patches along irrigation canals. Apparent reliance on vegetated patches, in an otherwise hostile human-dominated matrix, mirror findings that landscape heterogeneity and the presence of semi-natural vegetated features are required to maintain reptile diversity (Nopper et al. 2017, Pulsford et al. 2017, Boesing et al. 2018). The vulnerability of King Cobras in agricultural areas suggests that these areas may be acting as a population sink (Driscoll et al. 2013, Marshall et al. 2018), which emphasises the importance of maintaining vegetated areas within the landscape matrix to provide refuge from known mortality sources. Future research will assess spatial composition of resources available to King Cobras, and whether reduced movement leads to additional sub-lethal costs. More broadly, our findings suggest that wide-ranging reptiles can react to landscape fragmentation in similar ways to terrestrial mammals. This is especially important, because large snakes, such as King Cobras, fulfil underappreciated ecosystem roles (Miranda 2017). Their role in top-down trophic structuring is likely comparable to mammals that typically receive more conservation attention.

## Declarations

## Acknowledgements

We thank Nakhon Ratchasima Zoo, Dusit Zoo, and Zoological Park Organization under the Royal Patronage of His Majesty the King, Thailand; along with Wirongrong Changphet, Doctor of Veterinary Medicine (DVM); Wanlaya Tipkantha, DVM for their expertise in undertaking surgery on protected species. We thank the National Park, Wildlife and Plant Conservation Department, Thailand for supplying permits to study King Cobras. We thank the National Research Council of Thailand for providing permits for the project. We thank the Suranaree University of Technology and the School of Biology for supervising and funding the project, providing ethical approval, and general logistics. We thank Pluemjit Boonpueng for assisting with paperwork and logistics. We thank Assistant Professor Dr. Pantip Piyatadsananon, Vice director of Lower Northeast Regional Center of Geoinformatics and Space Technology Development Agency for obtaining land use data. We thank the Institute of Animals Scientific Purpose Development for supplying animal use licenses to C.T.S. and P.S. We thank Wildlife Reserves Singapore Conservation Fund, National Scientific and Technological Development Agency, and Herpetofauna Foundation for supplying funding and equipment. We thank the Thailand Institute of Scientific and Technological Research and Sakaerat Environmental Research Station for the consistent and crucial logistical support throughout the project. We thank the residents of Udom Sab for allowing research to be undertaken across their land. We thank the Hook 31 Rescue teams for their tireless work mitigating human-snake conflict and providing us with a number of King Cobras. We thank numerous Sakaerat Conservation and Snake Education Team members for countless hours tracking King Cobras throughout the landscape.

## Availability of data and materials

Data used in this study is available on Zenodo (DOI: 10.5281/zenodo.3666029) and Movebank (Movebank ID: 1093796277). The Zenodo repository also includes all R scripts used to run analysis and generate figures.

## Author contributions

*Conceptualization*, I.S., M.C., and C.T.S.; *Methodology*, I.S., M.C., C.T.S., and B.M.M.; *Formal Analysis*, B.M.M., M.C., and I.S.; *Investigation*, C.T.S., I.S., M.C., M.D.J., C.W.H. and B.M.M.; *Resources*, M.G., P.S., T.A., and S.W.; *Writing – Original Draft*, B.M.M., M.C., I.S., M.D.J., and C.T.S.; *Writing – Review & Editing*, B.M.M., M.C., I.S., M.D.J., C.W.H., C.T.S., and M.G.; *Visualisation*, B.M.M.; *Supervision*, P.S., S.W., and T.A.; *Funding Acquisition*, M.D.J., C.T.S., and P.S.

## Ethics approval and consent to participate

We had ethical approval from the Suranaree University of Technology Ethics Committee (24/2560). All work was undertaken Institute of Animals for Scientific Purpose Development (IAD) licenses belonging to P.S. and C.T.S. All work was permitted by the National Park, Wildlife and Plant Conservation Department, Thailand and the National Research Council of Thailand (98/59). All work was undertaken with permission from Thailand Institute of Scientific and Technological Research and Sakaerat Environmental Research Station.

