## Supplementary Figures and Tables for "No room to roam: King Cobras reduce movement in agriculture"

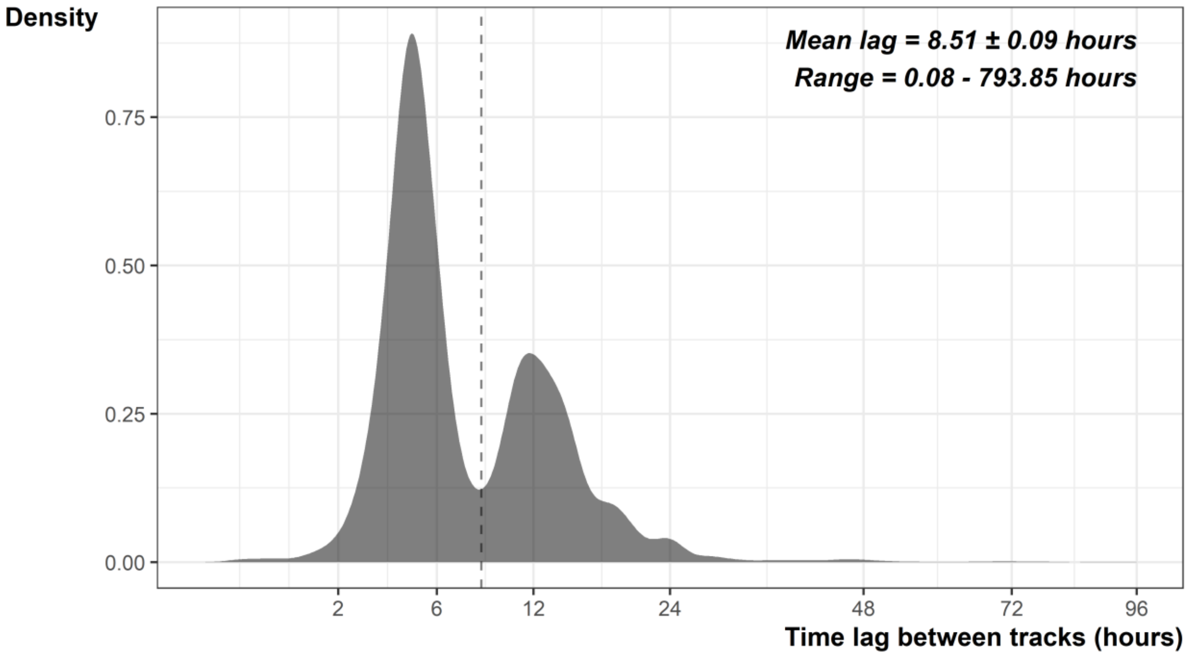


**Supplementary Figure 1. Distribution of time lags between radio tracking fixes.** Dashed lines indicate the mean time lag. X scale is log transformed and clipped at 96 hours for ease of visualisation.

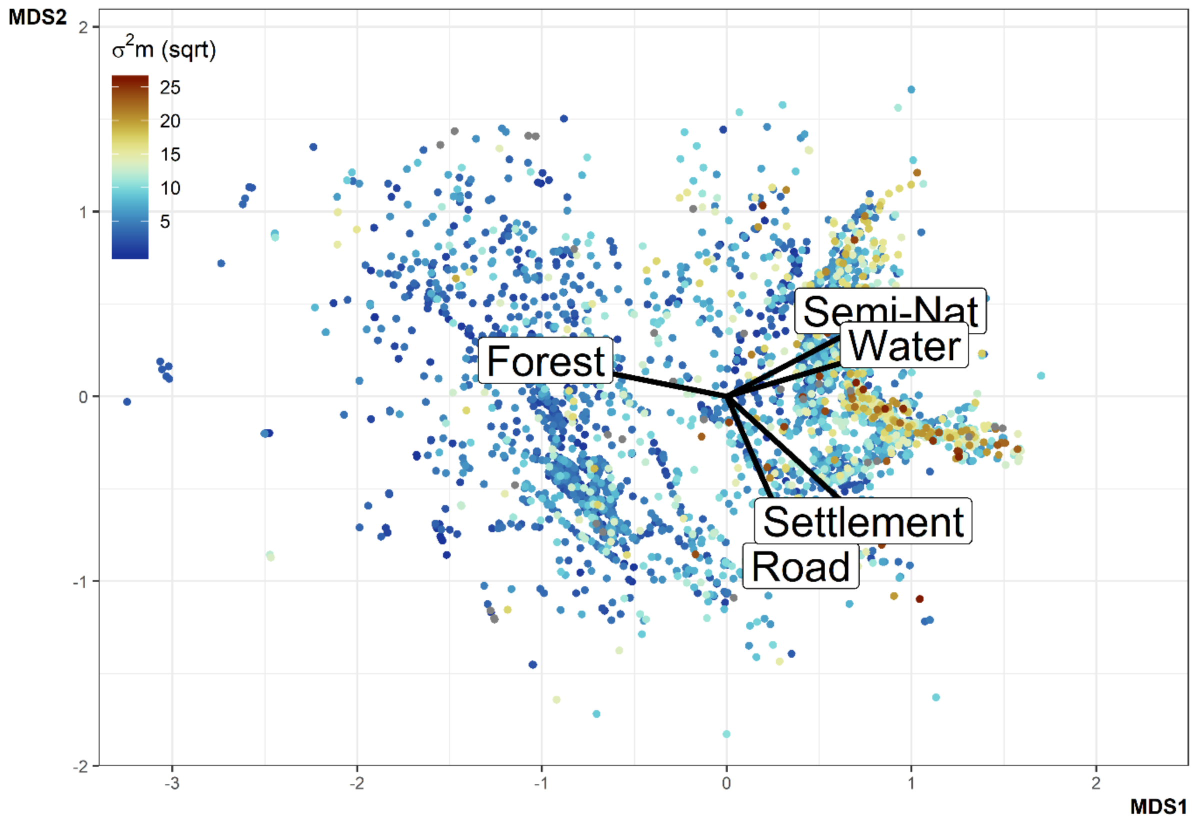


**Supplementary Figure 2. Bi-plot of NMDS results.** Motion variance values are reflected by the colour of the points, we have rooted these values so value differences are easier to distinguish.

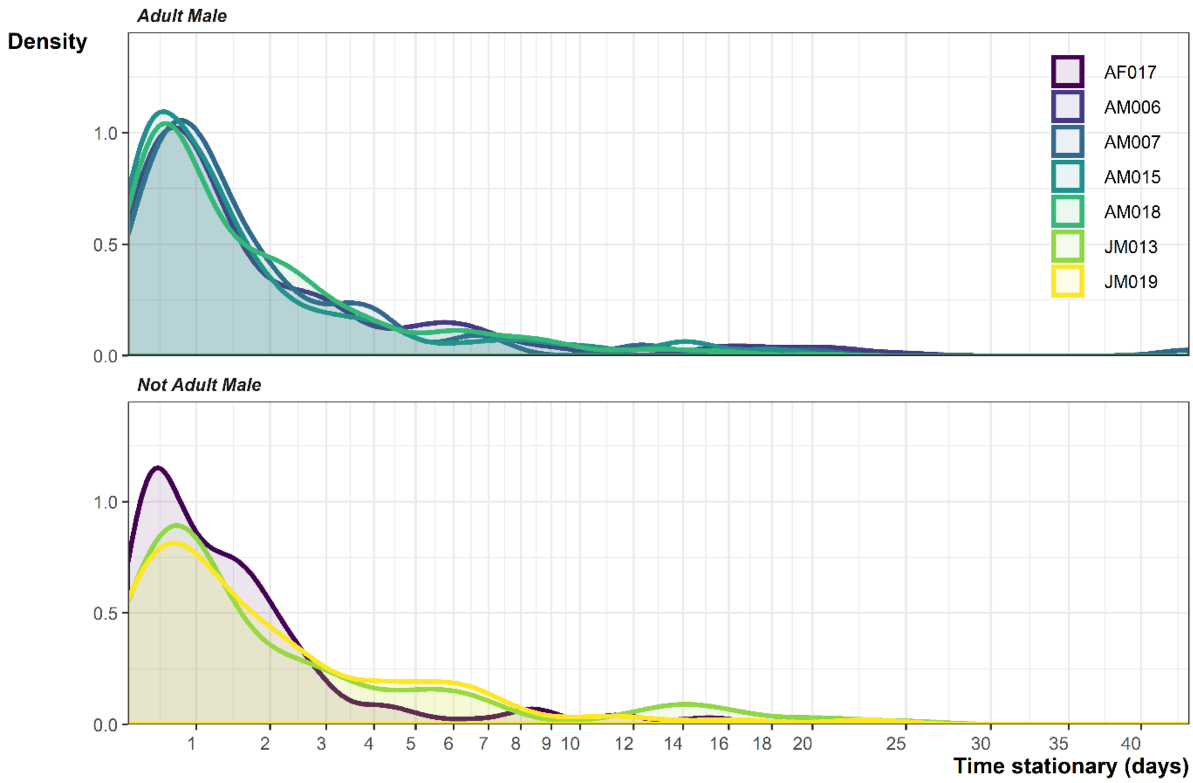


**Supplementary Figure 3. Distribution of sheltering periods.** To help distinguish individual lines the plots has been split in two. The top plot shows the results from the adult males: AM006, AM007, AM015 and AM018. The lower plot shows AF017, JM013 and JM019.

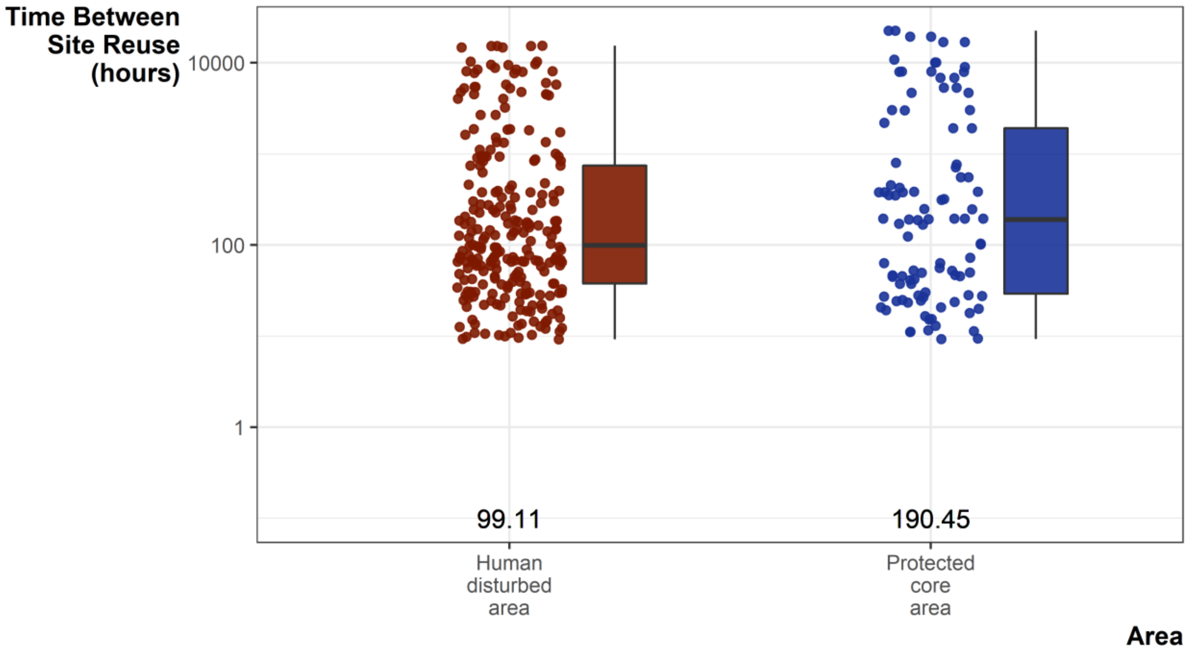


**Supplementary Figure 4. Bar charts for frequency of site reuse.** Labelled values are the median time between site reuse. Y axis is log transformed.

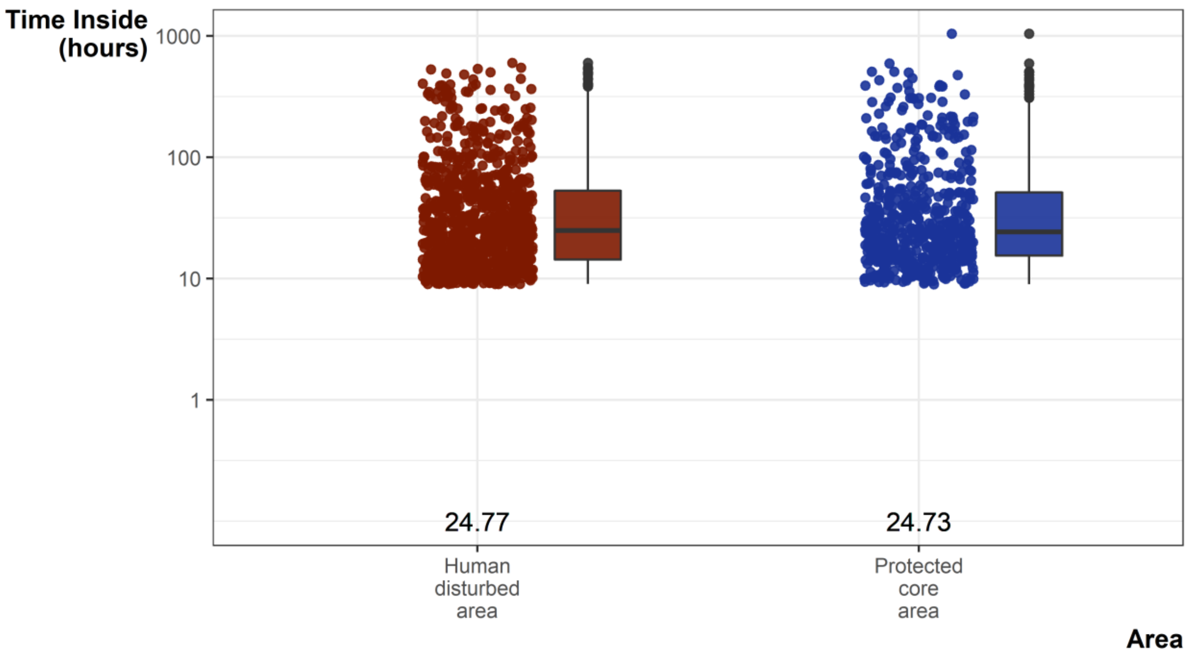


**Supplementary Figure 5. Bar charts for the sheltering times.** Labelled values are the median time between site reuse. Y axis is log transformed.

**Supplementary Table 1. All co-efficient results from Bayesian logistic regression models.**

| ID | Model | Parameter | Point Estimate | Lower CrI | Upper CrI |
|---|---|---|---|---|---|
| AF017 | residency_road+forest+semiNat | alpha | 1.199 | 1.165 | 1.235 |
| AM006 | residency_road+forest+semiNat | alpha | 1.204 | 1.170 | 1.232 |
| AM015 | residency_road+forest+semiNat | alpha | 1.203 | 1.172 | 1.232 |
| AM018 | residency_road+forest+semiNat | alpha | 1.213 | 1.180 | 1.248 |
| JM013 | residency_road+forest+semiNat | alpha | 1.226 | 1.187 | 1.276 |
| JM019 | residency_road+forest+semiNat | alpha | 1.221 | 1.188 | 1.282 |
| AF017 | residency_road+forest+semiNat | beta1 | 0.001 | -0.032 | 0.035 |
| AM006 | residency_road+forest+semiNat | beta1 | 0.005 | -0.017 | 0.028 |
| AM015 | residency_road+forest+semiNat | beta1 | -0.021 | -0.049 | 0.004 |
| AM018 | residency_road+forest+semiNat | beta1 | 0.002 | -0.020 | 0.028 |
| JM013 | residency_road+forest+semiNat | beta1 | 0.006 | -0.024 | 0.038 |
| JM019 | residency_road+forest+semiNat | beta1 | -0.018 | -0.048 | 0.009 |
| AF017 | residency_road+forest+semiNat | beta2 | 0.001 | -0.037 | 0.034 |
| AM006 | residency_road+forest+semiNat | beta2 | -0.014 | -0.057 | 0.020 |
| AM015 | residency_road+forest+semiNat | beta2 | 0.000 | -0.038 | 0.031 |
| AM018 | residency_road+forest+semiNat | beta2 | -0.011 | -0.059 | 0.026 |
| JM013 | residency_road+forest+semiNat | beta2 | 0.001 | -0.021 | 0.019 |
| JM019 | residency_road+forest+semiNat | beta2 | 0.008 | -0.021 | 0.039 |
| AF017 | residency_road+forest+semiNat | beta3 | 0.017 | -0.017 | 0.060 |
| AM006 | residency_road+forest+semiNat | beta3 | 0.026 | -0.005 | 0.056 |
| AM015 | residency_road+forest+semiNat | beta3 | -0.007 | -0.038 | 0.023 |
| AM018 | residency_road+forest+semiNat | beta3 | -0.012 | -0.039 | 0.006 |
| JM013 | residency_road+forest+semiNat | beta3 | -0.005 | -0.047 | 0.043 |

| | | | | | |
|---|---|---|---|---|---|
| JM019 | residency_road+forest+semiNat | beta3 | -0.010 | -0.056 | 0.036 |
| AF017 | residency_road+forest+settle | alpha | 1.186 | 1.159 | 1.210 |
| AM006 | residency_road+forest+settle | alpha | 1.205 | 1.174 | 1.240 |
| AM015 | residency_road+forest+settle | alpha | 1.200 | 1.173 | 1.226 |
| AM018 | residency_road+forest+settle | alpha | 1.206 | 1.179 | 1.231 |
| JM013 | residency_road+forest+settle | alpha | 1.228 | 1.189 | 1.278 |
| JM019 | residency_road+forest+settle | alpha | 1.224 | 1.185 | 1.266 |
| AF017 | residency_road+forest+settle | beta1 | 0.001 | -0.023 | 0.031 |
| AM006 | residency_road+forest+settle | beta1 | 0.004 | -0.016 | 0.024 |
| AM015 | residency_road+forest+settle | beta1 | -0.011 | -0.035 | 0.010 |
| AM018 | residency_road+forest+settle | beta1 | 0.000 | -0.018 | 0.023 |
| JM013 | residency_road+forest+settle | beta1 | 0.001 | -0.019 | 0.030 |
| JM019 | residency_road+forest+settle | beta1 | -0.011 | -0.036 | 0.011 |
| AF017 | residency_road+forest+settle | beta2 | 0.002 | -0.028 | 0.027 |
| AM006 | residency_road+forest+settle | beta2 | -0.008 | -0.044 | 0.021 |
| AM015 | residency_road+forest+settle | beta2 | 0.003 | -0.024 | 0.030 |
| AM018 | residency_road+forest+settle | beta2 | -0.002 | -0.037 | 0.025 |
| JM013 | residency_road+forest+settle | beta2 | 0.002 | -0.017 | 0.019 |
| JM019 | residency_road+forest+settle | beta2 | 0.006 | -0.016 | 0.032 |
| AF017 | residency_road+forest+settle | beta3 | -0.002 | -0.024 | 0.021 |
| AM006 | residency_road+forest+settle | beta3 | 0.008 | -0.012 | 0.031 |
| AM015 | residency_road+forest+settle | beta3 | -0.009 | -0.036 | 0.008 |
| AM018 | residency_road+forest+settle | beta3 | -0.012 | -0.034 | 0.005 |
| JM013 | residency_road+forest+settle | beta3 | -0.002 | -0.029 | 0.023 |
| JM019 | residency_road+forest+settle | beta3 | -0.007 | -0.042 | 0.017 |
| AF017 | revisit_road+forest+semiNat | alpha | 0.441 | 0.258 | 0.544 |
| AM006 | revisit_road+forest+semiNat | alpha | 0.143 | 0.011 | 0.260 |
| AM015 | revisit_road+forest+semiNat | alpha | 0.145 | 0.026 | 0.253 |
| AM018 | revisit_road+forest+semiNat | alpha | 0.333 | 0.206 | 0.454 |
| JM013 | revisit_road+forest+semiNat | alpha | 0.167 | 0.005 | 0.316 |
| JM019 | revisit_road+forest+semiNat | alpha | 0.317 | 0.160 | 0.486 |
| AF017 | revisit_road+forest+semiNat | beta1 | 0.007 | -0.081 | 0.087 |
| AM006 | revisit_road+forest+semiNat | beta1 | 0.017 | -0.042 | 0.090 |
| AM006 | revisit_road+forest+semiNat | beta1 | 0.017 | 0.103 | 0.103 |
| AM015 | revisit_road+forest+semiNat | beta1 | -0.003 | -0.091 | 0.065 |
| AM018 | revisit_road+forest+semiNat | beta1 | -0.005 | -0.094 | 0.050 |
| JM013 | revisit_road+forest+semiNat | beta1 | 0.006 | -0.081 | 0.078 |
| JM019 | revisit_road+forest+semiNat | beta1 | 0.021 | -0.039 | 0.104 |
| AF017 | revisit_road+forest+semiNat | beta2 | -0.021 | -0.160 | -0.152 |
| AF017 | revisit_road+forest+semiNat | beta2 | -0.021 | -0.136 | 0.049 |
| AM006 | revisit_road+forest+semiNat | beta2 | -0.003 | -0.079 | 0.109 |
| AM015 | revisit_road+forest+semiNat | beta2 | -0.015 | -0.114 | 0.075 |
| AM018 | revisit_road+forest+semiNat | beta2 | -0.011 | -0.124 | 0.076 |
| JM013 | revisit_road+forest+semiNat | beta2 | -0.005 | -0.055 | 0.056 |
| JM019 | revisit_road+forest+semiNat | beta2 | -0.027 | -0.136 | 0.037 |
| AF017 | revisit_road+forest+semiNat | beta3 | -0.010 | -0.183 | 0.067 |
| AM006 | revisit_road+forest+semiNat | beta3 | -0.009 | -0.112 | 0.049 |
| AM015 | revisit_road+forest+semiNat | beta3 | 0.000 | -0.090 | 0.077 |

| | | | | | |
|---|---|---|---|---|---|
| AM018 | revisit_road+forest+semiNat | beta3 | 0.013 | -0.041 | 0.084 |
| JM013 | revisit_road+forest+semiNat | beta3 | 0.008 | -0.088 | 0.121 |
| JM019 | revisit_road+forest+semiNat | beta3 | 0.016 | -0.066 | 0.142 |
| JM019 | revisit_road+forest+semiNat | beta3 | 0.016 | 0.145 | 0.157 |
| AF017 | revisit_road+forest+settle | alpha | 0.452 | 0.373 | 0.529 |
| AM006 | revisit_road+forest+settle | alpha | 0.138 | 0.013 | 0.251 |
| AM015 | revisit_road+forest+settle | alpha | 0.143 | 0.020 | 0.249 |
| AM018 | revisit_road+forest+settle | alpha | 0.358 | 0.273 | 0.443 |
| JM013 | revisit_road+forest+settle | alpha | 0.161 | 0.006 | 0.317 |
| JM019 | revisit_road+forest+settle | alpha | 0.318 | 0.170 | 0.469 |
| AF017 | revisit_road+forest+settle | beta1 | 0.006 | -0.062 | -0.055 |
| AF017 | revisit_road+forest+settle | beta1 | 0.006 | -0.048 | 0.071 |
| AM006 | revisit_road+forest+settle | beta1 | 0.013 | -0.034 | 0.075 |
| AM015 | revisit_road+forest+settle | beta1 | -0.001 | -0.053 | 0.057 |
| AM018 | revisit_road+forest+settle | beta1 | 0.000 | -0.059 | 0.047 |
| JM013 | revisit_road+forest+settle | beta1 | 0.005 | -0.070 | 0.055 |
| JM019 | revisit_road+forest+settle | beta1 | 0.011 | -0.041 | 0.074 |
| AF017 | revisit_road+forest+settle | beta2 | -0.013 | -0.093 | 0.037 |
| AM006 | revisit_road+forest+settle | beta2 | -0.005 | -0.075 | 0.072 |
| AM015 | revisit_road+forest+settle | beta2 | -0.008 | -0.094 | -0.084 |
| AM015 | revisit_road+forest+settle | beta2 | -0.008 | -0.077 | 0.054 |
| AM018 | revisit_road+forest+settle | beta2 | -0.008 | -0.078 | 0.065 |
| JM013 | revisit_road+forest+settle | beta2 | -0.003 | -0.051 | 0.047 |
| JM013 | revisit_road+forest+settle | beta2 | -0.003 | 0.050 | 0.054 |
| JM019 | revisit_road+forest+settle | beta2 | -0.015 | -0.097 | 0.032 |
| AF017 | revisit_road+forest+settle | beta3 | -0.001 | -0.091 | -0.091 |
| AF017 | revisit_road+forest+settle | beta3 | -0.001 | -0.072 | 0.052 |
| AM006 | revisit_road+forest+settle | beta3 | -0.001 | -0.068 | 0.041 |
| AM015 | revisit_road+forest+settle | beta3 | 0.002 | -0.054 | 0.055 |
| AM018 | revisit_road+forest+settle | beta3 | -0.003 | -0.067 | 0.035 |
| JM013 | revisit_road+forest+settle | beta3 | 0.003 | -0.060 | 0.061 |
| JM019 | revisit_road+forest+settle | beta3 | 0.016 | -0.032 | 0.109 |

**Supplementary Table 2. Full ISSF results for all models and individuals.**

| Term | Estimate | SE | Statistic | p-value | Conf.low | Conf.high | ID | Model | AIC |
|---|---|---|---|---|---|---|---|---|---|
| log_sl | 0.002 | 0.029 | 0.075 | 0.940 | -0.056 | 0.060 | AM006 | model1 | 5794.50 |
| cos_ta | -0.344 | 0.215 | -1.600 | 0.110 | -0.765 | 0.077 | AM006 | model1 | 5794.50 |
| log_sl:cos_ta | 0.068 | 0.042 | 1.642 | 0.101 | -0.013 | 0.150 | AM006 | model1 | 5794.50 |
| dist_forest | -2.598 | 2.784 | -0.933 | 0.351 | -8.054 | 2.858 | AM006 | model2 | 5794.62 |
| log_sl | -6.299 | 3.468 | -1.816 | 0.069 | -13.096 | 0.499 | AM006 | model2 | 5794.62 |
| cos_ta | -2.978 | 8.626 | -0.345 | 0.730 | -19.885 | 13.929 | AM006 | model2 | 5794.62 |
| dist_forest:log_sl | 0.801 | 0.441 | 1.817 | 0.069 | -0.063 | 1.665 | AM006 | model2 | 5794.62 |
| dist_forest:cos_ta | 0.332 | 1.097 | 0.303 | 0.762 | -1.817 | 2.481 | AM006 | model2 | 5794.62 |
| log_sl:cos_ta | 0.074 | 0.042 | 1.767 | 0.077 | -0.008 | 0.156 | AM006 | model2 | 5794.62 |
| dist_settle | 6.982 | 3.961 | 1.763 | 0.078 | -0.782 | 14.746 | AM006 | model3 | 5794.64 |
| log_sl | 10.479 | 5.290 | 1.981 | 0.048 | 0.110 | 20.848 | AM006 | model3 | 5794.64 |

| | | | | | | | | | |
|---|---|---|---|---|---|---|---|---|---|
| cos_ta | 13.572 | 10.547 | 1.287 | 0.198 | -7.099 | 34.243 | AM006 | model3 | 5794.64 |
| dist_settle:log_sl | -1.189 | 0.600 | -1.980 | 0.048 | -2.366 | -0.012 | AM006 | model3 | 5794.64 |
| dist_settle:cos_ta | -1.578 | 1.196 | -1.320 | 0.187 | -3.921 | 0.765 | AM006 | model3 | 5794.64 |
| log_sl:cos_ta | 0.065 | 0.042 | 1.571 | 0.116 | -0.016 | 0.147 | AM006 | model3 | 5794.64 |
| dist_semiNat | 5.887 | 1.654 | 3.558 | 0.000 | 2.645 | 9.130 | AM006 | model4 | 5783.74 |
| log_sl | 6.911 | 1.800 | 3.839 | 0.000 | 3.382 | 10.439 | AM006 | model4 | 5783.74 |
| cos_ta | 2.157 | 3.545 | 0.608 | 0.543 | -4.791 | 9.104 | AM006 | model4 | 5783.74 |
| dist_semiNat:log_sl | -0.787 | 0.205 | -3.843 | 0.000 | -1.188 | -0.386 | AM006 | model4 | 5783.74 |
| dist_semiNat:cos_ta | -0.280 | 0.400 | -0.699 | 0.485 | -1.065 | 0.505 | AM006 | model4 | 5783.74 |
| log_sl:cos_ta | 0.055 | 0.042 | 1.290 | 0.197 | -0.028 | 0.138 | AM006 | model4 | 5783.74 |
| dist_road | 3.627 | 1.767 | 2.053 | 0.040 | 0.164 | 7.090 | AM006 | model5 | 5791.24 |
| log_sl | 2.716 | 2.212 | 1.228 | 0.220 | -1.619 | 7.051 | AM006 | model5 | 5791.24 |
| cos_ta | 2.746 | 4.250 | 0.646 | 0.518 | -5.584 | 11.076 | AM006 | model5 | 5791.24 |
| dist_road:log_sl | -0.349 | 0.285 | -1.225 | 0.221 | -0.906 | 0.209 | AM006 | model5 | 5791.24 |
| dist_road:cos_ta | -0.396 | 0.545 | -0.727 | 0.467 | -1.463 | 0.672 | AM006 | model5 | 5791.24 |
| log_sl:cos_ta | 0.065 | 0.042 | 1.548 | 0.122 | -0.017 | 0.147 | AM006 | model5 | 5791.24 |
| dist_water | 3.242 | 1.332 | 2.434 | 0.015 | 0.632 | 5.853 | AM006 | model6 | 5779.88 |
| log_sl | 5.799 | 1.395 | 4.157 | 0.000 | 3.065 | 8.534 | AM006 | model6 | 5779.88 |
| cos_ta | 2.130 | 2.736 | 0.778 | 0.436 | -3.233 | 7.492 | AM006 | model6 | 5779.88 |
| dist_water:log_sl | -0.677 | 0.163 | -4.163 | 0.000 | -0.996 | -0.358 | AM006 | model6 | 5779.88 |
| dist_water:cos_ta | -0.291 | 0.316 | -0.921 | 0.357 | -0.909 | 0.328 | AM006 | model6 | 5779.88 |
| log_sl:cos_ta | 0.074 | 0.043 | 1.746 | 0.081 | -0.009 | 0.158 | AM006 | model6 | 5779.88 |
| dist_road | 1.642 | 0.607 | 2.704 | 0.007 | 0.452 | 2.833 | AM006 | model7 | 5789.37 |
| dist_forest | 2.311 | 1.339 | 1.725 | 0.085 | -0.315 | 4.936 | AM006 | model7 | 5789.37 |
| dist_semiNat | 1.080 | 1.087 | 0.994 | 0.320 | -1.049 | 3.210 | AM006 | model7 | 5789.37 |
| log_sl | 0.012 | 0.030 | 0.402 | 0.687 | -0.046 | 0.071 | AM006 | model7 | 5789.37 |
| cos_ta | -0.336 | 0.219 | -1.529 | 0.126 | -0.766 | 0.095 | AM006 | model7 | 5789.37 |
| log_sl:cos_ta | 0.065 | 0.043 | 1.496 | 0.135 | -0.020 | 0.150 | AM006 | model7 | 5789.37 |
| dist_road | 1.719 | 0.615 | 2.796 | 0.005 | 0.514 | 2.924 | AM006 | model8 | 5789.90 |
| dist_forest | 2.172 | 1.329 | 1.634 | 0.102 | -0.434 | 4.777 | AM006 | model8 | 5789.90 |
| dist_settle | -1.019 | 1.487 | -0.685 | 0.493 | -3.932 | 1.895 | AM006 | model8 | 5789.90 |
| log_sl | 0.012 | 0.030 | 0.386 | 0.699 | -0.047 | 0.070 | AM006 | model8 | 5789.90 |
| cos_ta | -0.372 | 0.217 | -1.711 | 0.087 | -0.798 | 0.054 | AM006 | model8 | 5789.90 |
| log_sl:cos_ta | 0.076 | 0.042 | 1.806 | 0.071 | -0.007 | 0.159 | AM006 | model8 | 5789.90 |
| dist_road | 1.593 | 0.609 | 2.617 | 0.009 | 0.400 | 2.786 | AM006 | model9 | 5789.75 |
| dist_forest | 2.091 | 1.335 | 1.566 | 0.117 | -0.526 | 4.708 | AM006 | model9 | 5789.75 |
| dist_water | -0.693 | 0.877 | -0.790 | 0.430 | -2.412 | 1.026 | AM006 | model9 | 5789.75 |
| log_sl | 0.012 | 0.030 | 0.387 | 0.699 | -0.047 | 0.070 | AM006 | model9 | 5789.75 |
| cos_ta | -0.391 | 0.220 | -1.779 | 0.075 | -0.821 | 0.040 | AM006 | model9 | 5789.75 |
| log_sl:cos_ta | 0.082 | 0.043 | 1.898 | 0.058 | -0.003 | 0.168 | AM006 | model9 | 5789.75 |
| log_sl | 0.002 | 0.026 | 0.096 | 0.923 | -0.048 | 0.053 | AM015 | model1 | 6205.87 |
| cos_ta | -0.390 | 0.180 | -2.164 | 0.030 | -0.743 | -0.037 | AM015 | model1 | 6205.87 |
| log_sl:cos_ta | 0.081 | 0.037 | 2.229 | 0.026 | 0.010 | 0.153 | AM015 | model1 | 6205.87 |
| dist_forest | -2.268 | 2.040 | -1.112 | 0.266 | -6.265 | 1.730 | AM015 | model2 | 6187.03 |
| log_sl | -8.003 | 2.498 | -3.204 | 0.001 | -12.899 | -3.107 | AM015 | model2 | 6187.03 |
| cos_ta | 5.667 | 6.159 | 0.920 | 0.358 | -6.404 | 17.738 | AM015 | model2 | 6187.03 |
| dist_forest:log_sl | 1.023 | 0.319 | 3.205 | 0.001 | 0.397 | 1.648 | AM015 | model2 | 6187.03 |
| dist_forest:cos_ta | -0.784 | 0.788 | -0.995 | 0.320 | -2.328 | 0.761 | AM015 | model2 | 6187.03 |

| | | | | | | | | | |
|---|---|---|---|---|---|---|---|---|---|
| log_sl:cos_ta | 0.110 | 0.037 | 2.956 | 0.003 | 0.037 | 0.182 | AM015 | model2 | 6187.03 |
| dist_settle | 12.938 | 4.199 | 3.081 | 0.002 | 4.708 | 21.169 | AM015 | model3 | 6201.81 |
| log_sl | 16.907 | 5.603 | 3.018 | 0.003 | 5.926 | 27.888 | AM015 | model3 | 6201.81 |
| cos_ta | -1.962 | 11.454 | -0.171 | 0.864 | -24.412 | 20.488 | AM015 | model3 | 6201.81 |
| dist_settle:log_sl | -1.914 | 0.634 | -3.018 | 0.003 | -3.156 | -0.671 | AM015 | model3 | 6201.81 |
| dist_settle:cos_ta | 0.178 | 1.295 | 0.138 | 0.890 | -2.359 | 2.716 | AM015 | model3 | 6201.81 |
| log_sl:cos_ta | 0.080 | 0.036 | 2.181 | 0.029 | 0.008 | 0.151 | AM015 | model3 | 6201.81 |
| dist_semiNat | 9.867 | 2.043 | 4.829 | 0.000 | 5.863 | 13.872 | AM015 | model4 | 6186.73 |
| log_sl | 4.790 | 2.413 | 1.985 | 0.047 | 0.060 | 9.519 | AM015 | model4 | 6186.73 |
| cos_ta | -6.860 | 5.069 | -1.353 | 0.176 | -16.795 | 3.076 | AM015 | model4 | 6186.73 |
| dist_semiNat:log_sl | -0.537 | 0.271 | -1.981 | 0.048 | -1.069 | -0.006 | AM015 | model4 | 6186.73 |
| dist_semiNat:cos_ta | 0.733 | 0.568 | 1.289 | 0.197 | -0.381 | 1.846 | AM015 | model4 | 6186.73 |
| log_sl:cos_ta | 0.063 | 0.037 | 1.677 | 0.094 | -0.011 | 0.136 | AM015 | model4 | 6186.73 |
| dist_road | 4.468 | 1.821 | 2.454 | 0.014 | 0.899 | 8.036 | AM015 | model5 | 6201.98 |
| log_sl | 6.556 | 2.211 | 2.965 | 0.003 | 2.223 | 10.890 | AM015 | model5 | 6201.98 |
| cos_ta | 0.520 | 4.453 | 0.117 | 0.907 | -8.208 | 9.247 | AM015 | model5 | 6201.98 |
| dist_road:log_sl | -0.844 | 0.285 | -2.967 | 0.003 | -1.402 | -0.286 | AM015 | model5 | 6201.98 |
| dist_road:cos_ta | -0.117 | 0.571 | -0.205 | 0.838 | -1.236 | 1.003 | AM015 | model5 | 6201.98 |
| log_sl:cos_ta | 0.081 | 0.036 | 2.233 | 0.026 | 0.010 | 0.153 | AM015 | model5 | 6201.98 |
| dist_water | 5.191 | 1.509 | 3.440 | 0.001 | 2.233 | 8.148 | AM015 | model6 | 6199.01 |
| log_sl | 4.162 | 1.602 | 2.599 | 0.009 | 1.023 | 7.301 | AM015 | model6 | 6199.01 |
| cos_ta | -4.428 | 3.341 | -1.325 | 0.185 | -10.976 | 2.120 | AM015 | model6 | 6199.01 |
| dist_water:log_sl | -0.479 | 0.184 | -2.599 | 0.009 | -0.840 | -0.118 | AM015 | model6 | 6199.01 |
| dist_water:cos_ta | 0.467 | 0.383 | 1.221 | 0.222 | -0.283 | 1.217 | AM015 | model6 | 6199.01 |
| log_sl:cos_ta | 0.073 | 0.037 | 1.960 | 0.050 | 0.000 | 0.145 | AM015 | model6 | 6199.01 |
| dist_road | 0.763 | 0.729 | 1.047 | 0.295 | -0.666 | 2.192 | AM015 | model7 | 6171.16 |
| dist_forest | 4.143 | 0.962 | 4.306 | 0.000 | 2.257 | 6.028 | AM015 | model7 | 6171.16 |
| dist_semiNat | 8.340 | 1.634 | 5.104 | 0.000 | 5.138 | 11.542 | AM015 | model7 | 6171.16 |
| log_sl | 0.023 | 0.027 | 0.884 | 0.377 | -0.029 | 0.075 | AM015 | model7 | 6171.16 |
| cos_ta | -0.388 | 0.185 | -2.095 | 0.036 | -0.751 | -0.025 | AM015 | model7 | 6171.16 |
| log_sl:cos_ta | 0.080 | 0.038 | 2.091 | 0.037 | 0.005 | 0.155 | AM015 | model7 | 6171.16 |
| dist_road | -0.642 | 0.776 | -0.827 | 0.408 | -2.163 | 0.879 | AM015 | model8 | 6195.03 |
| dist_forest | 3.296 | 0.911 | 3.618 | 0.000 | 1.511 | 5.082 | AM015 | model8 | 6195.03 |
| dist_settle | 2.600 | 1.993 | 1.305 | 0.192 | -1.306 | 6.507 | AM015 | model8 | 6195.03 |
| log_sl | 0.011 | 0.026 | 0.438 | 0.662 | -0.040 | 0.063 | AM015 | model8 | 6195.03 |
| cos_ta | -0.450 | 0.182 | -2.468 | 0.014 | -0.808 | -0.093 | AM015 | model8 | 6195.03 |
| log_sl:cos_ta | 0.103 | 0.037 | 2.758 | 0.006 | 0.030 | 0.177 | AM015 | model8 | 6195.03 |
| dist_road | -0.043 | 0.695 | -0.062 | 0.950 | -1.406 | 1.319 | AM015 | model9 | 6187.41 |
| dist_forest | 3.947 | 0.952 | 4.144 | 0.000 | 2.080 | 5.814 | AM015 | model9 | 6187.41 |
| dist_water | 3.606 | 1.178 | 3.062 | 0.002 | 1.298 | 5.914 | AM015 | model9 | 6187.41 |
| log_sl | 0.016 | 0.026 | 0.589 | 0.556 | -0.036 | 0.067 | AM015 | model9 | 6187.41 |
| cos_ta | -0.411 | 0.184 | -2.236 | 0.025 | -0.772 | -0.051 | AM015 | model9 | 6187.41 |
| log_sl:cos_ta | 0.090 | 0.038 | 2.359 | 0.018 | 0.015 | 0.164 | AM015 | model9 | 6187.41 |
| log_sl | 0.001 | 0.025 | 0.038 | 0.970 | -0.048 | 0.050 | AF017 | model1 | 7573.47 |
| cos_ta | -0.347 | 0.147 | -2.358 | 0.018 | -0.635 | -0.059 | AF017 | model1 | 7573.47 |
| log_sl:cos_ta | 0.082 | 0.035 | 2.320 | 0.020 | 0.013 | 0.151 | AF017 | model1 | 7573.47 |
| dist_forest | 6.749 | 2.642 | 2.554 | 0.011 | 1.570 | 11.928 | AF017 | model2 | 7555.47 |
| log_sl | 2.564 | 3.675 | 0.698 | 0.485 | -4.638 | 9.766 | AF017 | model2 | 7555.47 |

| | | | | | | | | | |
|---|---|---|---|---|---|---|---|---|---|
| cos_ta | -6.687 | 7.388 | -0.905 | 0.365 | -21.168 | 7.794 | AF017 | model2 | 7555.47 |
| dist_forest:log_sl | -0.327 | 0.471 | -0.694 | 0.488 | -1.249 | 0.596 | AF017 | model2 | 7555.47 |
| dist_forest:cos_ta | 0.804 | 0.946 | 0.850 | 0.395 | -1.049 | 2.657 | AF017 | model2 | 7555.47 |
| log_sl:cos_ta | 0.113 | 0.036 | 3.134 | 0.002 | 0.043 | 0.184 | AF017 | model2 | 7555.47 |
| dist_settle | -11.608 | 6.486 | -1.790 | 0.073 | -24.321 | 1.104 | AF017 | model3 | 7542.05 |
| log_sl | 15.698 | 9.068 | 1.731 | 0.083 | -2.075 | 33.471 | AF017 | model3 | 7542.05 |
| cos_ta | 17.120 | 17.916 | 0.956 | 0.339 | -17.994 | 52.234 | AF017 | model3 | 7542.05 |
| dist_settle:log_sl | -1.772 | 1.025 | -1.730 | 0.084 | -3.780 | 0.236 | AF017 | model3 | 7542.05 |
| dist_settle:cos_ta | -1.979 | 2.024 | -0.978 | 0.328 | -5.946 | 1.988 | AF017 | model3 | 7542.05 |
| log_sl:cos_ta | 0.110 | 0.036 | 3.063 | 0.002 | 0.039 | 0.180 | AF017 | model3 | 7542.05 |
| dist_semiNat | 49.638 | 6.039 | 8.219 | 0.000 | 37.801 | 61.475 | AF017 | model4 | 7492.52 |
| log_sl | 5.493 | 9.832 | 0.559 | 0.576 | -13.778 | 24.763 | AF017 | model4 | 7492.52 |
| cos_ta | -22.353 | 19.731 | -1.133 | 0.257 | -61.025 | 16.319 | AF017 | model4 | 7492.52 |
| dist_semiNat:log_sl | -0.609 | 1.098 | -0.555 | 0.579 | -2.760 | 1.543 | AF017 | model4 | 7492.52 |
| dist_semiNat:cos_ta | 2.458 | 2.202 | 1.116 | 0.264 | -1.857 | 6.773 | AF017 | model4 | 7492.52 |
| log_sl:cos_ta | 0.078 | 0.036 | 2.148 | 0.032 | 0.007 | 0.150 | AF017 | model4 | 7492.52 |
| dist_road | 3.042 | 3.462 | 0.879 | 0.380 | -3.744 | 9.828 | AF017 | model5 | 7557.50 |
| log_sl | 12.230 | 4.620 | 2.647 | 0.008 | 3.175 | 21.284 | AF017 | model5 | 7557.50 |
| cos_ta | 4.479 | 8.592 | 0.521 | 0.602 | -12.362 | 21.320 | AF017 | model5 | 7557.50 |
| dist_road:log_sl | -1.566 | 0.592 | -2.646 | 0.008 | -2.725 | -0.406 | AF017 | model5 | 7557.50 |
| dist_road:cos_ta | -0.621 | 1.100 | -0.565 | 0.572 | -2.776 | 1.534 | AF017 | model5 | 7557.50 |
| log_sl:cos_ta | 0.093 | 0.035 | 2.634 | 0.008 | 0.024 | 0.163 | AF017 | model5 | 7557.50 |
| dist_water | 13.119 | 4.678 | 2.804 | 0.005 | 3.950 | 22.288 | AF017 | model6 | 7566.34 |
| log_sl | 19.036 | 6.296 | 3.024 | 0.002 | 6.696 | 31.376 | AF017 | model6 | 7566.34 |
| cos_ta | 14.500 | 10.794 | 1.343 | 0.179 | -6.657 | 35.657 | AF017 | model6 | 7566.34 |
| dist_water:log_sl | -2.172 | 0.718 | -3.024 | 0.002 | -3.580 | -0.764 | AF017 | model6 | 7566.34 |
| dist_water:cos_ta | -1.692 | 1.231 | -1.375 | 0.169 | -4.104 | 0.720 | AF017 | model6 | 7566.34 |
| log_sl:cos_ta | 0.073 | 0.036 | 2.049 | 0.040 | 0.003 | 0.143 | AF017 | model6 | 7566.34 |
| dist_road | -0.652 | 1.678 | -0.389 | 0.698 | -3.940 | 2.637 | AF017 | model7 | 7471.48 |
| dist_forest | 5.103 | 1.326 | 3.847 | 0.000 | 2.503 | 7.703 | AF017 | model7 | 7471.48 |
| dist_semiNat | 46.099 | 4.885 | 9.437 | 0.000 | 36.525 | 55.673 | AF017 | model7 | 7471.48 |
| log_sl | 0.052 | 0.026 | 1.948 | 0.051 | 0.000 | 0.103 | AF017 | model7 | 7471.48 |
| cos_ta | -0.400 | 0.153 | -2.612 | 0.009 | -0.699 | -0.100 | AF017 | model7 | 7471.48 |
| log_sl:cos_ta | 0.110 | 0.037 | 2.929 | 0.003 | 0.036 | 0.183 | AF017 | model7 | 7471.48 |
| dist_road | -1.496 | 1.627 | -0.920 | 0.358 | -4.684 | 1.693 | AF017 | model8 | 7539.43 |
| dist_forest | 2.511 | 1.294 | 1.940 | 0.052 | -0.025 | 5.048 | AF017 | model8 | 7539.43 |
| dist_settle | -16.100 | 4.265 | -3.775 | 0.000 | -24.459 | -7.741 | AF017 | model8 | 7539.43 |
| log_sl | 0.019 | 0.025 | 0.750 | 0.453 | -0.031 | 0.069 | AF017 | model8 | 7539.43 |
| cos_ta | -0.430 | 0.149 | -2.880 | 0.004 | -0.722 | -0.137 | AF017 | model8 | 7539.43 |
| log_sl:cos_ta | 0.125 | 0.036 | 3.443 | 0.001 | 0.054 | 0.197 | AF017 | model8 | 7539.43 |
| dist_road | -2.641 | 1.635 | -1.616 | 0.106 | -5.846 | 0.563 | AF017 | model9 | 7530.33 |
| dist_forest | 8.751 | 1.593 | 5.495 | 0.000 | 5.630 | 11.872 | AF017 | model9 | 7530.33 |
| dist_water | 16.967 | 3.491 | 4.860 | 0.000 | 10.124 | 23.810 | AF017 | model9 | 7530.33 |
| log_sl | 0.027 | 0.026 | 1.059 | 0.290 | -0.023 | 0.078 | AF017 | model9 | 7530.33 |
| cos_ta | -0.398 | 0.151 | -2.641 | 0.008 | -0.693 | -0.103 | AF017 | model9 | 7530.33 |
| log_sl:cos_ta | 0.105 | 0.037 | 2.858 | 0.004 | 0.033 | 0.178 | AF017 | model9 | 7530.33 |
| log_sl | -0.006 | 0.040 | -0.146 | 0.884 | -0.085 | 0.073 | JM013 | model1 | 4014.21 |
| cos_ta | -0.289 | 0.277 | -1.043 | 0.297 | -0.831 | 0.254 | JM013 | model1 | 4014.21 |

| | | | | | | | | | |
|---|---|---|---|---|---|---|---|---|---|
| log_sl:cos_ta | 0.057 | 0.057 | 0.993 | 0.321 | -0.055 | 0.169 | JM013 | model1 | 4014.21 |
| dist_forest | 0.109 | 1.122 | 0.097 | 0.923 | -2.091 | 2.308 | JM013 | model2 | 4013.58 |
| log_sl | -0.721 | 1.243 | -0.580 | 0.562 | -3.157 | 1.715 | JM013 | model2 | 4013.58 |
| cos_ta | -4.996 | 2.311 | -2.162 | 0.031 | -9.525 | -0.467 | JM013 | model2 | 4013.58 |
| dist_forest:log_sl | 0.095 | 0.163 | 0.580 | 0.562 | -0.225 | 0.414 | JM013 | model2 | 4013.58 |
| dist_forest:cos_ta | 0.614 | 0.302 | 2.031 | 0.042 | 0.022 | 1.206 | JM013 | model2 | 4013.58 |
| log_sl:cos_ta | 0.067 | 0.059 | 1.141 | 0.254 | -0.048 | 0.181 | JM013 | model2 | 4013.58 |
| dist_settle | 3.421 | 5.699 | 0.600 | 0.548 | -7.749 | 14.591 | JM013 | model3 | 4014.60 |
| log_sl | 4.489 | 8.144 | 0.551 | 0.581 | -11.472 | 20.451 | JM013 | model3 | 4014.60 |
| cos_ta | 32.701 | 14.761 | 2.215 | 0.027 | 3.769 | 61.633 | JM013 | model3 | 4014.60 |
| dist_settle:log_sl | -0.508 | 0.921 | -0.552 | 0.581 | -2.313 | 1.297 | JM013 | model3 | 4014.60 |
| dist_settle:cos_ta | -3.730 | 1.669 | -2.235 | 0.025 | -7.000 | -0.460 | JM013 | model3 | 4014.60 |
| log_sl:cos_ta | 0.052 | 0.058 | 0.907 | 0.364 | -0.061 | 0.165 | JM013 | model3 | 4014.60 |
| dist_semiNat | -0.565 | 4.585 | -0.123 | 0.902 | -9.552 | 8.423 | JM013 | model4 | 4017.76 |
| log_sl | -1.083 | 6.436 | -0.168 | 0.866 | -13.697 | 11.532 | JM013 | model4 | 4017.76 |
| cos_ta | 18.186 | 11.936 | 1.524 | 0.128 | -5.208 | 41.580 | JM013 | model4 | 4017.76 |
| dist_semiNat:log_sl | 0.122 | 0.725 | 0.168 | 0.867 | -1.299 | 1.542 | JM013 | model4 | 4017.76 |
| dist_semiNat:cos_ta | -2.080 | 1.344 | -1.548 | 0.122 | -4.713 | 0.553 | JM013 | model4 | 4017.76 |
| log_sl:cos_ta | 0.056 | 0.058 | 0.976 | 0.329 | -0.057 | 0.170 | JM013 | model4 | 4017.76 |
| dist_road | -0.205 | 3.231 | -0.064 | 0.949 | -6.537 | 6.127 | JM013 | model5 | 4017.31 |
| log_sl | 0.660 | 4.421 | 0.149 | 0.881 | -8.004 | 9.324 | JM013 | model5 | 4017.31 |
| cos_ta | 11.256 | 7.639 | 1.473 | 0.141 | -3.717 | 26.229 | JM013 | model5 | 4017.31 |
| dist_road:log_sl | -0.085 | 0.568 | -0.150 | 0.881 | -1.199 | 1.029 | JM013 | model5 | 4017.31 |
| dist_road:cos_ta | -1.485 | 0.982 | -1.513 | 0.130 | -3.409 | 0.439 | JM013 | model5 | 4017.31 |
| log_sl:cos_ta | 0.058 | 0.057 | 1.007 | 0.314 | -0.055 | 0.170 | JM013 | model5 | 4017.31 |
| dist_water | 15.882 | 6.458 | 2.459 | 0.014 | 3.225 | 28.539 | JM013 | model6 | 4013.23 |
| log_sl | 19.349 | 9.910 | 1.953 | 0.051 | -0.073 | 38.772 | JM013 | model6 | 4013.23 |
| cos_ta | -12.201 | 17.538 | -0.696 | 0.487 | -46.574 | 22.173 | JM013 | model6 | 4013.23 |
| dist_water:log_sl | -2.204 | 1.128 | -1.953 | 0.051 | -4.415 | 0.007 | JM013 | model6 | 4013.23 |
| dist_water:cos_ta | 1.355 | 1.994 | 0.680 | 0.497 | -2.553 | 5.263 | JM013 | model6 | 4013.23 |
| log_sl:cos_ta | 0.059 | 0.057 | 1.030 | 0.303 | -0.053 | 0.171 | JM013 | model6 | 4013.23 |
| dist_road | -0.647 | 1.040 | -0.622 | 0.534 | -2.685 | 1.391 | JM013 | model7 | 4017.55 |
| dist_forest | 1.011 | 0.708 | 1.427 | 0.154 | -0.378 | 2.399 | JM013 | model7 | 4017.55 |
| dist_semiNat | 1.877 | 2.773 | 0.677 | 0.498 | -3.558 | 7.312 | JM013 | model7 | 4017.55 |
| log_sl | -0.004 | 0.040 | -0.100 | 0.921 | -0.083 | 0.075 | JM013 | model7 | 4017.55 |
| cos_ta | -0.330 | 0.279 | -1.182 | 0.237 | -0.877 | 0.217 | JM013 | model7 | 4017.55 |
| log_sl:cos_ta | 0.070 | 0.058 | 1.202 | 0.229 | -0.044 | 0.185 | JM013 | model7 | 4017.55 |
| dist_road | -0.509 | 1.034 | -0.492 | 0.622 | -2.535 | 1.517 | JM013 | model8 | 4017.61 |
| dist_forest | 0.983 | 0.697 | 1.410 | 0.159 | -0.384 | 2.350 | JM013 | model8 | 4017.61 |
| dist_settle | 1.914 | 3.028 | 0.632 | 0.527 | -4.021 | 7.849 | JM013 | model8 | 4017.61 |
| log_sl | -0.005 | 0.040 | -0.113 | 0.910 | -0.083 | 0.074 | JM013 | model8 | 4017.61 |
| cos_ta | -0.334 | 0.279 | -1.200 | 0.230 | -0.881 | 0.212 | JM013 | model8 | 4017.61 |
| log_sl:cos_ta | 0.072 | 0.058 | 1.234 | 0.217 | -0.042 | 0.186 | JM013 | model8 | 4017.61 |
| dist_road | -0.624 | 1.034 | -0.604 | 0.546 | -2.652 | 1.403 | JM013 | model9 | 4015.71 |
| dist_forest | 0.704 | 0.646 | 1.090 | 0.276 | -0.562 | 1.971 | JM013 | model9 | 4015.71 |
| dist_water | 4.254 | 2.806 | 1.516 | 0.130 | -1.247 | 9.755 | JM013 | model9 | 4015.71 |
| log_sl | -0.003 | 0.040 | -0.079 | 0.937 | -0.082 | 0.076 | JM013 | model9 | 4015.71 |
| cos_ta | -0.326 | 0.279 | -1.168 | 0.243 | -0.873 | 0.221 | JM013 | model9 | 4015.71 |

| | | | | | | | | | |
|---|---|---|---|---|---|---|---|---|---|
| log_sl:cos_ta | 0.069 | 0.058 | 1.181 | 0.238 | -0.045 | 0.183 | JM013 | model9 | 4015.71 |
| log_sl | -0.001 | 0.041 | -0.036 | 0.971 | -0.082 | 0.079 | JM019 | model1 | 2402.41 |
| cos_ta | -0.209 | 0.269 | -0.776 | 0.437 | -0.737 | 0.319 | JM019 | model1 | 2402.41 |
| log_sl:cos_ta | 0.048 | 0.058 | 0.826 | 0.409 | -0.066 | 0.163 | JM019 | model1 | 2402.41 |
| dist_forest | 1.300 | 1.837 | 0.708 | 0.479 | -2.299 | 4.900 | JM019 | model2 | 2399.25 |
| log_sl | -3.101 | 2.029 | -1.528 | 0.127 | -7.079 | 0.877 | JM019 | model2 | 2399.25 |
| cos_ta | -5.265 | 5.065 | -1.039 | 0.299 | -15.192 | 4.662 | JM019 | model2 | 2399.25 |
| dist_forest:log_sl | 0.398 | 0.261 | 1.529 | 0.126 | -0.112 | 0.909 | JM019 | model2 | 2399.25 |
| dist_forest:cos_ta | 0.649 | 0.650 | 0.998 | 0.318 | -0.625 | 1.922 | JM019 | model2 | 2399.25 |
| log_sl:cos_ta | 0.046 | 0.059 | 0.773 | 0.440 | -0.070 | 0.162 | JM019 | model2 | 2399.25 |
| dist_settle | 11.810 | 5.934 | 1.990 | 0.047 | 0.179 | 23.441 | JM019 | model3 | 2402.45 |
| log_sl | 14.968 | 7.738 | 1.934 | 0.053 | -0.198 | 30.134 | JM019 | model3 | 2402.45 |
| cos_ta | 19.452 | 16.343 | 1.190 | 0.234 | -12.581 | 51.484 | JM019 | model3 | 2402.45 |
| dist_settle:log_sl | -1.695 | 0.876 | -1.935 | 0.053 | -3.413 | 0.022 | JM019 | model3 | 2402.45 |
| dist_settle:cos_ta | -2.223 | 1.848 | -1.203 | 0.229 | -5.846 | 1.399 | JM019 | model3 | 2402.45 |
| log_sl:cos_ta | 0.034 | 0.059 | 0.579 | 0.562 | -0.082 | 0.150 | JM019 | model3 | 2402.45 |
| dist_semiNat | 17.157 | 4.575 | 3.750 | 0.000 | 8.190 | 26.123 | JM019 | model4 | 2392.32 |
| log_sl | 10.508 | 6.093 | 1.725 | 0.085 | -1.434 | 22.451 | JM019 | model4 | 2392.32 |
| cos_ta | 10.952 | 12.849 | 0.852 | 0.394 | -14.231 | 36.135 | JM019 | model4 | 2392.32 |
| dist_semiNat:log_sl | -1.179 | 0.684 | -1.724 | 0.085 | -2.519 | 0.162 | JM019 | model4 | 2392.32 |
| dist_semiNat:cos_ta | -1.251 | 1.439 | -0.869 | 0.385 | -4.071 | 1.569 | JM019 | model4 | 2392.32 |
| log_sl:cos_ta | 0.042 | 0.059 | 0.708 | 0.479 | -0.074 | 0.158 | JM019 | model4 | 2392.32 |
| dist_road | 1.191 | 1.989 | 0.599 | 0.549 | -2.708 | 5.090 | JM019 | model5 | 2406.31 |
| log_sl | 2.312 | 2.248 | 1.028 | 0.304 | -2.095 | 6.719 | JM019 | model5 | 2406.31 |
| cos_ta | 3.119 | 4.759 | 0.656 | 0.512 | -6.208 | 12.446 | JM019 | model5 | 2406.31 |
| dist_road:log_sl | -0.300 | 0.292 | -1.029 | 0.303 | -0.872 | 0.272 | JM019 | model5 | 2406.31 |
| dist_road:cos_ta | -0.433 | 0.615 | -0.704 | 0.481 | -1.638 | 0.772 | JM019 | model5 | 2406.31 |
| log_sl:cos_ta | 0.051 | 0.059 | 0.862 | 0.389 | -0.065 | 0.166 | JM019 | model5 | 2406.31 |
| dist_water | 6.567 | 3.990 | 1.646 | 0.100 | -1.252 | 14.387 | JM019 | model6 | 2404.11 |
| log_sl | 6.300 | 4.933 | 1.277 | 0.202 | -3.368 | 15.968 | JM019 | model6 | 2404.11 |
| cos_ta | 13.353 | 11.228 | 1.189 | 0.234 | -8.655 | 35.360 | JM019 | model6 | 2404.11 |
| dist_water:log_sl | -0.721 | 0.565 | -1.277 | 0.201 | -1.828 | 0.385 | JM019 | model6 | 2404.11 |
| dist_water:cos_ta | -1.550 | 1.284 | -1.208 | 0.227 | -4.066 | 0.966 | JM019 | model6 | 2404.11 |
| log_sl:cos_ta | 0.038 | 0.059 | 0.651 | 0.515 | -0.077 | 0.153 | JM019 | model6 | 2404.11 |
| dist_road | -1.449 | 1.080 | -1.342 | 0.180 | -3.567 | 0.668 | JM019 | model7 | 2388.28 |
| dist_forest | 3.354 | 1.381 | 2.428 | 0.015 | 0.647 | 6.061 | JM019 | model7 | 2388.28 |
| dist_semiNat | 12.804 | 3.405 | 3.760 | 0.000 | 6.130 | 19.477 | JM019 | model7 | 2388.28 |
| log_sl | 0.020 | 0.042 | 0.472 | 0.637 | -0.063 | 0.102 | JM019 | model7 | 2388.28 |
| cos_ta | -0.252 | 0.275 | -0.917 | 0.359 | -0.791 | 0.287 | JM019 | model7 | 2388.28 |
| log_sl:cos_ta | 0.065 | 0.061 | 1.078 | 0.281 | -0.053 | 0.184 | JM019 | model7 | 2388.28 |
| dist_road | -2.259 | 1.465 | -1.543 | 0.123 | -5.130 | 0.611 | JM019 | model8 | 2400.04 |
| dist_forest | 3.162 | 1.349 | 2.344 | 0.019 | 0.518 | 5.807 | JM019 | model8 | 2400.04 |
| dist_settle | 7.222 | 4.797 | 1.506 | 0.132 | -2.180 | 16.625 | JM019 | model8 | 2400.04 |
| log_sl | 0.009 | 0.042 | 0.209 | 0.834 | -0.073 | 0.090 | JM019 | model8 | 2400.04 |
| cos_ta | -0.210 | 0.272 | -0.774 | 0.439 | -0.744 | 0.323 | JM019 | model8 | 2400.04 |
| log_sl:cos_ta | 0.048 | 0.060 | 0.801 | 0.423 | -0.069 | 0.165 | JM019 | model8 | 2400.04 |
| dist_road | -1.771 | 1.075 | -1.648 | 0.099 | -3.877 | 0.335 | JM019 | model9 | 2398.12 |
| dist_forest | 3.758 | 1.402 | 2.680 | 0.007 | 1.010 | 6.506 | JM019 | model9 | 2398.12 |

| | | | | | | | | | |
|---|---|---|---|---|---|---|---|---|---|
| dist_water | 6.688 | 3.187 | 2.098 | 0.036 | 0.441 | 12.934 | JM019 | model9 | 2398.12 |
| log_sl | 0.009 | 0.042 | 0.208 | 0.835 | -0.073 | 0.090 | JM019 | model9 | 2398.12 |
| cos_ta | -0.220 | 0.271 | -0.811 | 0.417 | -0.751 | 0.311 | JM019 | model9 | 2398.12 |
| log_sl:cos_ta | 0.053 | 0.059 | 0.890 | 0.374 | -0.064 | 0.169 | JM019 | model9 | 2398.12 |
| log_sl | -0.001 | 0.017 | -0.049 | 0.961 | -0.035 | 0.033 | AM018 | model1 | 10567.77 |
| cos_ta | -0.179 | 0.118 | -1.517 | 0.129 | -0.410 | 0.052 | AM018 | model1 | 10567.77 |
| log_sl:cos_ta | 0.037 | 0.024 | 1.530 | 0.126 | -0.010 | 0.085 | AM018 | model1 | 10567.77 |
| dist_forest | 0.203 | 4.978 | 0.041 | 0.968 | -9.554 | 9.959 | AM018 | model2 | 10530.19 |
| log_sl | -19.517 | 6.817 | -2.863 | 0.004 | -32.878 | -6.156 | AM018 | model2 | 10530.19 |
| cos_ta | 30.593 | 22.751 | 1.345 | 0.179 | -13.998 | 75.184 | AM018 | model2 | 10530.19 |
| dist_forest:log_sl | 2.475 | 0.864 | 2.864 | 0.004 | 0.781 | 4.169 | AM018 | model2 | 10530.19 |
| dist_forest:cos_ta | -3.905 | 2.884 | -1.354 | 0.176 | -9.557 | 1.748 | AM018 | model2 | 10530.19 |
| log_sl:cos_ta | 0.050 | 0.025 | 2.028 | 0.043 | 0.002 | 0.099 | AM018 | model2 | 10530.19 |
| dist_settle | 3.389 | 2.610 | 1.299 | 0.194 | -1.726 | 8.505 | AM018 | model3 | 10552.30 |
| log_sl | -3.141 | 3.823 | -0.822 | 0.411 | -10.634 | 4.352 | AM018 | model3 | 10552.30 |
| cos_ta | -18.552 | 10.014 | -1.853 | 0.064 | -38.180 | 1.076 | AM018 | model3 | 10552.30 |
| dist_settle:log_sl | 0.356 | 0.433 | 0.823 | 0.411 | -0.492 | 1.205 | AM018 | model3 | 10552.30 |
| dist_settle:cos_ta | 2.081 | 1.134 | 1.835 | 0.066 | -0.141 | 4.303 | AM018 | model3 | 10552.30 |
| log_sl:cos_ta | 0.035 | 0.025 | 1.437 | 0.151 | -0.013 | 0.083 | AM018 | model3 | 10552.30 |
| dist_semiNat | 0.915 | 1.117 | 0.819 | 0.413 | -1.275 | 3.105 | AM018 | model4 | 10563.71 |
| log_sl | 2.765 | 1.095 | 2.526 | 0.012 | 0.619 | 4.910 | AM018 | model4 | 10563.71 |
| cos_ta | 2.237 | 2.743 | 0.815 | 0.415 | -3.139 | 7.612 | AM018 | model4 | 10563.71 |
| dist_semiNat:log_sl | -0.319 | 0.126 | -2.528 | 0.011 | -0.566 | -0.072 | AM018 | model4 | 10563.71 |
| dist_semiNat:cos_ta | -0.280 | 0.315 | -0.887 | 0.375 | -0.898 | 0.338 | AM018 | model4 | 10563.71 |
| log_sl:cos_ta | 0.042 | 0.025 | 1.690 | 0.091 | -0.007 | 0.090 | AM018 | model4 | 10563.71 |
| dist_road | 5.911 | 1.508 | 3.920 | 0.000 | 2.955 | 8.867 | AM018 | model5 | 10498.78 |
| log_sl | 1.082 | 2.065 | 0.524 | 0.600 | -2.966 | 5.130 | AM018 | model5 | 10498.78 |
| cos_ta | -6.882 | 4.964 | -1.386 | 0.166 | -16.611 | 2.847 | AM018 | model5 | 10498.78 |
| dist_road:log_sl | -0.135 | 0.264 | -0.512 | 0.608 | -0.653 | 0.382 | AM018 | model5 | 10498.78 |
| dist_road:cos_ta | 0.855 | 0.633 | 1.351 | 0.177 | -0.386 | 2.096 | AM018 | model5 | 10498.78 |
| log_sl:cos_ta | 0.042 | 0.025 | 1.673 | 0.094 | -0.007 | 0.092 | AM018 | model5 | 10498.78 |
| dist_water | 2.708 | 1.305 | 2.075 | 0.038 | 0.150 | 5.267 | AM018 | model6 | 10559.55 |
| log_sl | 4.721 | 1.437 | 3.286 | 0.001 | 1.905 | 7.537 | AM018 | model6 | 10559.55 |
| cos_ta | 2.857 | 3.416 | 0.836 | 0.403 | -3.838 | 9.552 | AM018 | model6 | 10559.55 |
| dist_water:log_sl | -0.551 | 0.168 | -3.289 | 0.001 | -0.879 | -0.223 | AM018 | model6 | 10559.55 |
| dist_water:cos_ta | -0.355 | 0.397 | -0.893 | 0.372 | -1.134 | 0.424 | AM018 | model6 | 10559.55 |
| log_sl:cos_ta | 0.040 | 0.025 | 1.616 | 0.106 | -0.008 | 0.088 | AM018 | model6 | 10559.55 |
| dist_road | 5.118 | 0.653 | 7.834 | 0.000 | 3.838 | 6.399 | AM018 | model7 | 10471.26 |
| dist_forest | 9.739 | 2.360 | 4.127 | 0.000 | 5.114 | 14.365 | AM018 | model7 | 10471.26 |
| dist_semiNat | -1.743 | 0.831 | -2.098 | 0.036 | -3.371 | -0.115 | AM018 | model7 | 10471.26 |
| log_sl | 0.034 | 0.018 | 1.840 | 0.066 | -0.002 | 0.069 | AM018 | model7 | 10471.26 |
| cos_ta | -0.228 | 0.123 | -1.854 | 0.064 | -0.469 | 0.013 | AM018 | model7 | 10471.26 |
| log_sl:cos_ta | 0.057 | 0.026 | 2.217 | 0.027 | 0.007 | 0.108 | AM018 | model7 | 10471.26 |
| dist_road | 4.515 | 0.677 | 6.672 | 0.000 | 3.189 | 5.841 | AM018 | model8 | 10472.37 |
| dist_forest | 10.353 | 2.358 | 4.391 | 0.000 | 5.732 | 14.975 | AM018 | model8 | 10472.37 |
| dist_settle | 2.504 | 1.402 | 1.786 | 0.074 | -0.244 | 5.253 | AM018 | model8 | 10472.37 |
| log_sl | 0.034 | 0.018 | 1.871 | 0.061 | -0.002 | 0.070 | AM018 | model8 | 10472.37 |
| cos_ta | -0.207 | 0.123 | -1.688 | 0.091 | -0.448 | 0.033 | AM018 | model8 | 10472.37 |

| | | | | | | | | |
|---|---|---|---|---|---|---|---|---|
| log_sl:cos_ta | 0.048 | 0.026 | 1.868 | 0.062 | -0.002 | 0.098 | AM018 | model8 | 10472.37 |
| dist_road | 4.981 | 0.647 | 7.699 | 0.000 | 3.713 | 6.248 | AM018 | model9 | 10474.02 |
| dist_forest | 9.954 | 2.369 | 4.201 | 0.000 | 5.310 | 14.598 | AM018 | model9 | 10474.02 |
| dist_water | -0.966 | 0.759 | -1.274 | 0.203 | -2.453 | 0.521 | AM018 | model9 | 10474.02 |
| log_sl | 0.033 | 0.018 | 1.801 | 0.072 | -0.003 | 0.068 | AM018 | model9 | 10474.02 |
| cos_ta | -0.220 | 0.123 | -1.789 | 0.074 | -0.461 | 0.021 | AM018 | model9 | 10474.02 |
| log_sl:cos_ta | 0.053 | 0.026 | 2.073 | 0.038 | 0.003 | 0.104 | AM018 | model9 | 10474.02 |
